# Microarray Gene Expression Dataset Re-Analysis Reveals Variability in Influenza Infection and Vaccination

**DOI:** 10.1101/702068

**Authors:** Lavida R. K. Rogers, Gustavo de los Campos, George I. Mias

## Abstract

Influenza, a communicable disease, affects thousands of people worldwide. Young children, elderly, immunocompromised individuals and pregnant women are at higher risk for being infected by the influenza virus. Our study aims to highlight differentially expressed genes in influenza disease compared to influenza vaccination. We also investigate genetic variation due to the age and sex of samples. To accomplish our goals, we conducted a meta-analysis using publicly available microarray expression data. Our inclusion criteria included subjects with influenza, subjects who received the influenza vaccine and healthy controls. We curated 18 microarray datasets for a total of 3,481 samples (1,277 controls, 297 influenza infection, 1,907 influenza vaccination). We pre-processed the raw microarray expression data in R using packages available to pre-process Affymetrix and Illumina microarray platforms. We used a Box-Cox power transformation of the data prior to our down-stream analysis to identify differentially expressed genes. Statistical analyses were based on linear mixed effects model with all study factors and successive likelihood ratio tests (LRT) to identify differentially-expressed genes. We filtered LRT results by disease (Bonferroni adjusted p-value *<* 0.05) and used a two-tailed 10% quantile cutoff to identify biologically significant genes. Furthermore, we assessed age and sex effects on the disease genes by filtering for genes with a statistically significant (Bonferroni adjusted p-value *<* 0.05) interaction between disease and age, and disease and sex. We identified 4,889 statistically significant genes when we filtered the LRT results by disease factor, and gene enrichment analysis (gene ontology and pathways) included innate immune response, viral process, defense response to virus, Hematopoietic cell lineage and NF-kappa B signaling pathway. Our quantile filtered gene lists comprised of 978 genes each associated with influenza infection and vaccination. We also identified 907 and 48 genes with statistically significant (Bonferroni adjusted p-value *<* 0.05) disease-age and disease-sex interactions respectively. Our meta-analysis approach highlights key gene signatures and their associated pathways for both influenza infection and vaccination. We also were able to identify genes with an age and sex effect. This gives potential for improving current vaccines and exploring genes that are expressed equally across ages when considering universal vaccinations for influenza.

## 1 INTRODUCTION

The influenza virus, a respiratory pathogen, is responsible for seasonal influenza (also known as the flu), influenza pandemics and high rates of morbidity and mortality worldwide (1). The influenza virus infects the upper respiratory tract by invading the epithelial cells, releasing viral RNA, replicating and spreading throughout the respiratory tract while also causing inflammation (2). Influenza is a highly contagious disease and spreads easily via contact with an infected person’s nasal discharges and cough droplets (3). The main virulence factors are haemagglutinin (HA) and neuraminidase (NA) (2). These surface glycoproteins are also important for determining the sub-type of the influenza virus. The influenza virus can also reduce host gene expression through their viral proteins (4, 5). The viral proteins affect transcription and translation in the host which reduces the production of host proteins and promotes immune system evasion for the virus (4, 5). The virus interferes with host gene expression to promote viral gene expression, and this affects the immune system of the host by reducing the expression of immune components such as the major histocompatibility (MHC) molecules antigen presentation, and interferon and cytokine signaling pathways (4, 6).

Influenza is a global health burden, and as a preventative method vaccinations are offered annually. Vaccines are modified annually because the influenza virus strains change and mutate every season (7). The influenza vaccinations target the viral strains and sub-types that researchers predict would be most prevalent each flu season (8, 3). Furthermore, there are groups in the population who are considered at a higher risk for influenza infection, and they include young children, elderly, individuals who are immunocompromised, and females who are pregnant (3). The Centers for Disease Control and Prevention (CDC) has estimated, for the 2017-2018 season for influenza, 959,000 hospitalizations and over 79,000 deaths (3). 90% of the deaths during the 2017-2018 flu season were within the elderly population, while about 48,000 of the hospitalizations were in children (3). These estimates highlight that young children and especially the elderly are at higher risks for influenza and severe infections that can lead to hospitalization or death. Additionally, the CDC has recommended varying dosages for each vaccine for different age groups due to age-dependent immune responses (9, 3). Due to a decrease in efficacy of the influenza vaccines in the 65 and older population, they receive different dosages compared to younger age groups, in order to elicit a beneficial immune response (9, 3). Contrasting between changes in gene expression due to immunosenescence in healthy subjects and the age-dependent immune responses to diseases such as influenza can help our understanding of how responses to different diseases vary with age. Due to the influenza virus constantly changing and the efficacy of the vaccine being dependent on one’s age, researchers have started efforts to develop a universal vaccine (10, 11, 12). The goal is for such a universal vaccine to provide protection to all influenza strains (13). One approach, is to implement the use of highly conserved influenza peptides in vaccine formulations (12, 13).

Previous studies have investigated global blood gene expression to compare influenza disease to other respiratory diseases to assess severity and pathogenesis (14). For example, influenza has been shown to induce a stronger immune response than respiratory syncytial virus by producing more respiratory cytokines (14, 15). Studies also explored responses to vaccinations to highlight gene signatures. In our meta-analysis, our aim was to combine publicly available influenza microarray data to identify the effects of disease state (control, influenza infection and vaccination), age and sex on gene expression. We explored gene expression variation in blood for 3,481 samples (1,277 controls, 297 influenza infected, 1,907 influenza vaccinated) to identify genes and their pathways in influenza (Figure 1-2). This is to the best of our knowledge, the largest meta-analysis (18 datasets) to explore blood expression changes in influenza infection and vaccination. Our results provide gene signatures and pathways that can be targeted to improve influenza treatment and vaccinations. We also highlight disease associated genes that have interactions with age and sex, that can be used to further explore improving vaccinations, and aid efforts in identifying potential gene targets towards developing universal vaccinations to help reduce the burden of influenza.

**Figure 1.**
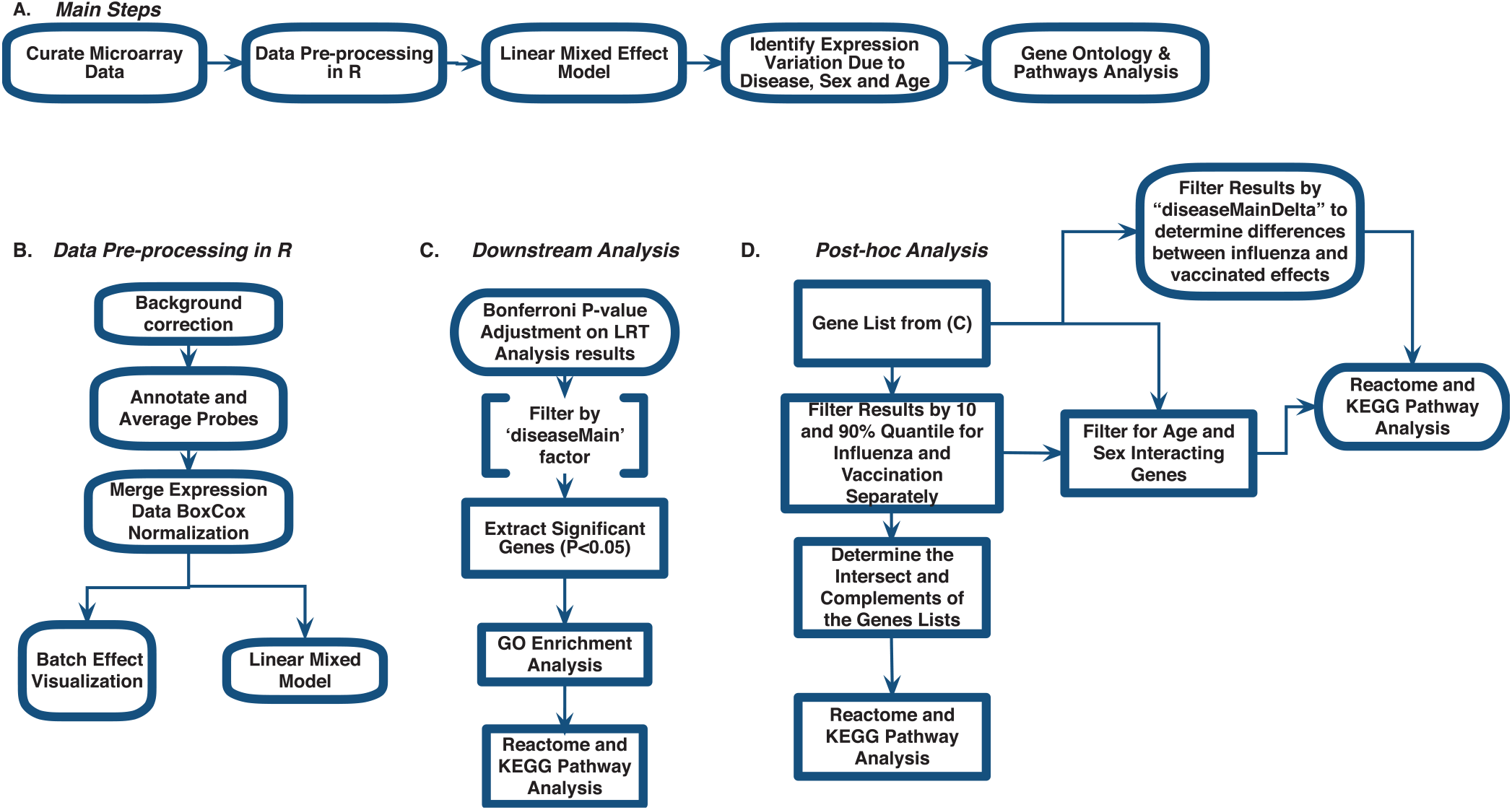
Meta-analysis Workflow to Assess Gene Expression Variation in Influenza Disease and Vaccination

## 2 METHODS

We curated 18 influenza-related microarray datasets from public database repositories (Table 1) to investigate changes in gene expression due to disease status, sex and age. The 18 datasets were from Affymetrix and Illumina microarray platforms (Table 1). We modified and implemented the data-analysis pipeline outlined by Brooks et al.(16)). To achieve our goal, after curating the datasets, we used the R programming language (17) to pre-process the raw gene expression data and to fit linear mixed effects models to determine statistically significant differentially expressed genes by factor (Figure 1). In addition, we identified genes that varied in expression due to disease status, sex, and age, and we also determined which gene ontology (GO) terms and pathways enrichment based on these gene sets (Figure 1).

**Table 1.** Demographics of curated influenza microarray datasets.

| Accession Number | Controls | Influenza Disease | Influenza Vaccine | Sex (M/F) | Age Range | Platform | Ref |
| --- | --- | --- | --- | --- | --- | --- | --- |
| GSE38900 | 31 | 16 | 0 | 20/27 | 0.025 - 1.57 | Illumina HumanHT-12 V4.0 expression beadchip | (14) |
| GSE107990 | 171 | 0 | 500 | 238/433 | 23 - 89 | Illumina HumanHT-12 V4.0 expression beadchip | (57) |
| GSE111368 | 130 | 229 | 0 | 177/182 | 18 - 71 | Illumina HumanHT-12 V4.0 expression beadchip | (46) |
| GSE27131 | 7 | 14 | 0 | 16/5 | 25 - 59 | Affymetrix Human Gene 1.0 ST Array | (58) |
| GSE29614 | 9 | 0 | 18 | 12/15 | 22 - 46 | Affymetrix Human Genome U133 Plus 2.0 Array | (53) |
| GSE29615 | 28 | 0 | 55 | 38/45 | 21 - 46 | Affymetrix HT HG-U133+ PM Array Plate | (53) |
| GSE47353 | 117 | 0 | 175 | 122/170 | 21 - 62 | Affymetrix Human Gene 1.0 ST Array | (59) |
| GSE48762 | 274 | 0 | 150 | 202/222 | 22 - 49 | Illumina HumanHT-12 V3.0 expression beadchip | (60) |
| GSE50628 | 0 | 10 | 0 | 2/8 | 4 - 9 | Affymetrix Human Genome U133 Plus 2.0 Array | (61) |
| GSE52005 | 34 | 0 | 102 | 62/74 | 0.68 - 14.68 | Illumina HumanHT-12 V4.0 expression beadchip | (62) |
| GSE74816 | 72 | 0 | 105 | 59/118 | 21 - 80 | Affymetrix HT HG-U133+ PM Array Plate | (55) |
| GSE97485 | 10 | 0 | 0 | 6/4 | 27 - 72 | Affymetrix Human Gene 1.0 ST Array | (63) |
| GSE34205 | 18 | 28 | 0 | 24/22 | 0.0416 - 11 | Affymetrix Human Genome U133 Plus 2.0 Array | (15) |
| GSE41080 | 91 | 0 | 0 | 37/54 | 20 - 93 | Illumina HumanHT-12 V3.0 expression beadchip | (64) |
| GSE74811 | 28 | 0 | 55 | 23/60 | 21 - 47 | Affymetrix HT HG-U133+ PM Array Plate | (55) |
| GSE59654 | 39 | 0 | 117 | 68/88 | 22 - 90 | Illumina HumanHT-12 V4.0 expression beadchip | (65) |
| GSE48018 | 111 | 0 | 320 | 431/0 | 18.2 - 32.1 | Illumina HumanHT-12 V3.0 expression beadchip | (66) |
| GSE48023 | 107 | 0 | 310 | 0/417 | 18.5-40.2 | Illumina HumanHT-12 V4.0 expression beadchip | (66) |

### 2.1 Data Curation: Gene Expression Omnibus

For our meta-analysis, we focused on influenza infection and vaccination. We searched public database repositories such as Gene Expression Omnibus (GEO) (18), Array Express (AE) (19) and Immune Space (IS) (20, 21) (Figure 2). To begin our data search, we found datasets with the keyword “influenza” and filtered for /textitHomo sapiens (Figure 2). Following this filter, we then removed duplicate records. For example, there were 15 duplicate records on GEO and 16 datasets on IS overlapped with our GEO records (Figure 2). We further filtered the results for datasets that were published, had non-ambiguous annotation, reported the age and sex of all subjects, and used blood or peripheral blood mononuclear cells (PBMCs) as the tissue type (Figure 2). Based on our inclusion criteria, we identified 18 datasets on GEO to use for our meta-analysis (Table 1 and SDF1 of online supplementary data files). For datasets such as GSE29614 (SDY64 on IS), GSE29615 (SDY269 on IS), GSE74811 (SDY270 on IS), GSE59654 (SDY404 on IS), GSE74816 (SDY1119 on IS), GSE48023 (SDY1276 on IS), 48018 (SDY1276 on IS) that did not have the ages of the subjects reported on GEO, we used the annotation from IS to gather age and sex characteristics of the samples. Additionally, we excluded 4 duplicates in GSE34205: GSM844139, GSM844141, GSM844143 and GSM844196 (which are duplicates of GSM844138, GSM844140, GSM844142, and GSM844195 datasets respectively).

**Figure 2.**
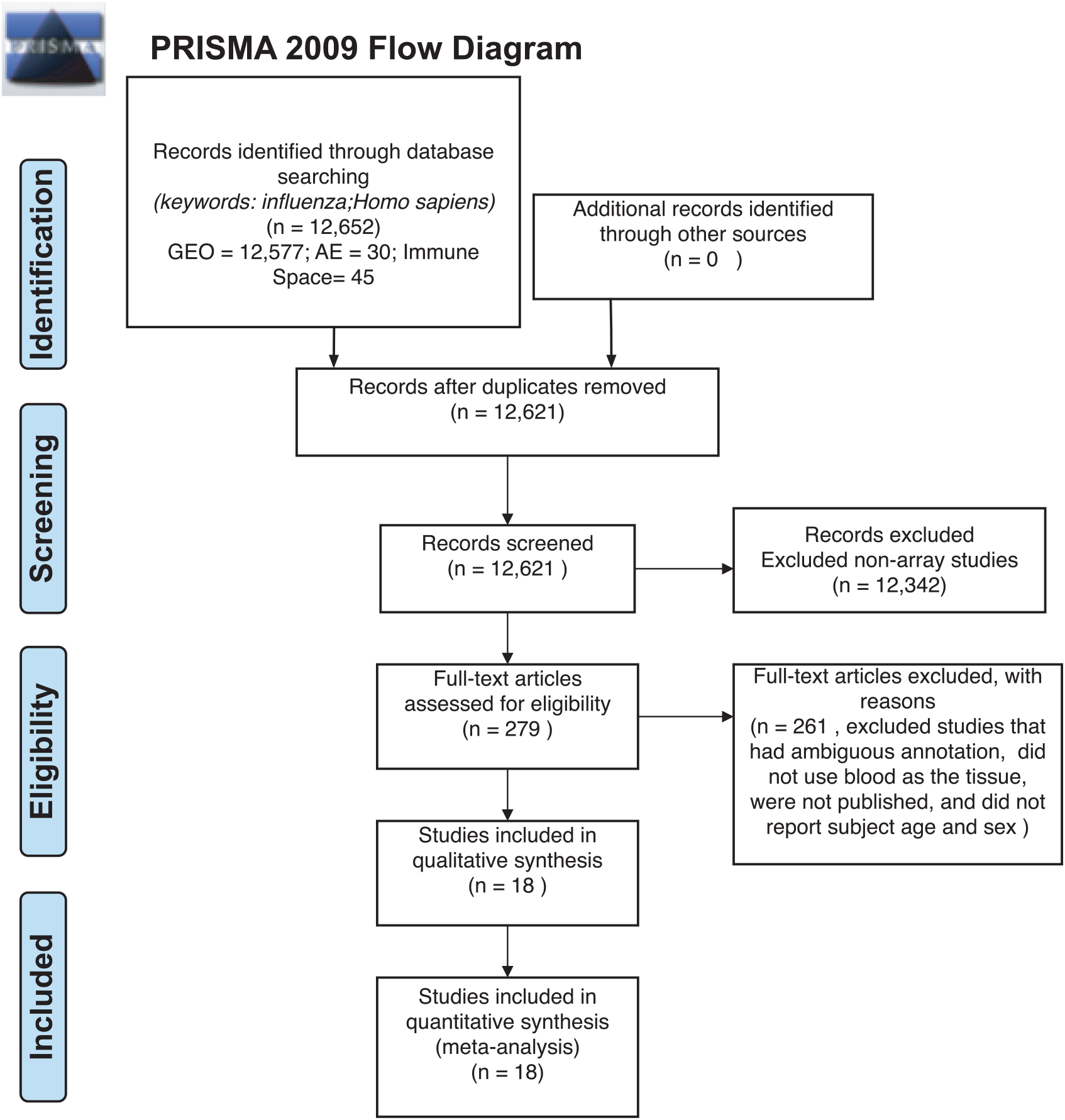
Preferred Reporting Items for Systematic Reviews and Meta-Analyses (PRISMA) Checklist.

After filtering through and selecting the datasets to use in our meta-analysis, we downloaded the raw gene expression data for each dataset, and created a file per study with sample characteristics (Table 1 and SDF1 of online supplementary data files). Our selected datasets were further filtered to remove samples that did not fit our criteria. For instance, GSE38900 and GSE34205 have samples with respiratory syncytial virus (RSV), GSE48762 contains samples who received the pneumococcal vaccine, GSE50628 has samples with rota-virus infection and patients who experience seizures, and GSE97485 has samples with acute myeloid leukemia who received the influenza vaccine. Due to this, we excluded all subjects that had a pre-existing health condition, infections other than influenza and received vaccinations other than the influenza vaccine (SDF1 of online supplementary data files).

### 2.2 Data Pre-Processing in R and Mathematica

All raw expression files were downloaded directly from the GEO website and pre-processed in R using appropriate packages based on the type of microarray platform (Table 1). We carried out background correction and annotated and summarized all probes (Fig 1B). We used the `affy` package (22) to pre-process all of the data files for the expression data from Affymetrix Human Genome Plus 2.0 and the Affymetrix HT Human Genome U133 Plus PM. Specifically, we used the `expresso` function to pre-process the files using robust multi-array analysis (RMA) for background correction, conduct perfect-match probe correction, and to calculate expression values using ‘avdiff’ (22). To summarize and remove replicate probes we used the `avereps` function from `limma` (23). For the Affymetrix HT Human Genome U133 Plus PM, we created our own annotation package in R using the annotation obtained from GEO (17). For the raw expression data from the Affymetrix Human Gene 1.1 ST microarray platform, we pre-processed the data using the `oligo` (24) and `affycoretools` (25) packages. To background correct the Affymetrix Human Gene 1.1 ST microarray data files we also used RMA and summarized and removed replicate probes using `avereps` function from `limma`. Our Illumina data files were pre-processed with the `limma` package. We used the NormExp Background Correction (`nec`) function from the limma package to remove the background of data files that reported the detection p-values. The (`nec`) function using the detection p-values when background correcting. Probes were annotated and summarized using the `aggregate` function from the `stats` package in base R (23, 17).

Following pre-processing, we merged expression data for the 18 datasets (Table 1 and SDF1 of online supplementary data files) by matching gene symbols that were common across all datasets. We conducted a Box-Cox power transformation (26) and standardized the expression values using the functions `ApplyBoxCoxTransformExtended` and `StandardizeExtended` from the MathIOmica (version 1.2.0) package in Mathematica (27, 28) (Fig 1B and SDF2 of online supplementary data files).

### 2.3 Linear Mixed Effects Modeling

We fitted a sequence of mixed-effects models to identify genes whose expression levels were affected by disease status (3 levels: control, influenza, vaccine) and those for which the effect of disease was modulated by either age or sex. Models were fitted using the lmer function od the `lme4` R-package (29). Separate models were fitted to each of the genes. Our baseline model (M0) included the (fixed) effects of sex (M/F), age (a factor with 4 levels, (−1,3],(3,19], (19,65] and (65,100]), ethnicity (a factor with 7 levels, African-American, Caucasian, Asian, Hispanic, Middle Eastern, Other, Unclassified) and tissue (2 levels, blood and PBMCs) plus the random effects of study (18 levels, see Table 1 for accession numbers) and of the subject (we included the subject effect because some studies had repeated measures). We first expanded this model by adding the (fixed) main effect of disease status (a factor with three levels, M1). Our next model expanded M1 by adding interactions between disease status and age (M2-DxA) and disease status by sex (M2-DxS). P-values for the main effects of diseases as for disease-by-sex and disease-by-age were obtained using likelihood ratio tests (LRT) between the models described above (SDF3 of online supplementary data files). LRTs were implemented using the anova function from base R to pairs of models. We used a sequential testing approach whereas: (i) we first identified genes with significant main effect of disease (this was based on a LRT between M1 and M0), (ii) among genes with significant main effect of diseases we tested the significance of DxA and DxS using a likelihood ratio test that had M1 as null hypothesis and the interaction models as alternative hypotheses. P-values were adjusted using Bonferroni, where for the first test (i) the number of tests was equal to the number of genes, and for the second one (ii) the number of tests was equal to the number of genes that passed the first test.

The filtering of genes based on Bonferroni-adjusted p-values for the main effect of disease (comparison of M1 to M0) allowed us to identify differentially expressed genes with respect to disease states (Figure 1). Using this gene list, we then conducted GO enrichment analysis (`GOAnalysis` function in `MathIOmica` package) and pathway enrichment analysis using Kyoto Encyclopedia of Genes and Genomes (KEGG, using the `KEGGAnalysis` functions in `MathIOmica`), and Reactome pathway enrichment analysis (`enrichPathway` function from the `ReactomePA` package in R (30)).

### 2.4 Determining Gene Expression Variability between Influenza Infection and Vaccination

We took a sequential testing approach to further analyze the identified statistically significant disease genes (SDF5 of online supplementary data files). Using this gene list, we further filtered for biological effect by using calculated estimates (which compared influenza and vaccine expression to controls) (SDF4 of online supplementary data files) and performed a two-tailed 10% quantile filter (i.e. 0.1 and 0.9 quantiles) to determine genes that were biologically significant in subjects who were vaccinated with influenza vaccinated and subjects infected with influenza disease. The biologically significant gene lists for the vaccinated and influenza subjects were further examined to identify genes in common, and genes only in the influenza list, and only in the vaccinated list (Figure 1). We performed GO and pathway enrichment analysis on these genes. Lastly, we filtered the disease (see SDF1 of online supplementary data files) statistically significant gene list for interacting genes between disease and age (age groups: (−1,3],(3,19], (19,65], (65,100])) and disease and sex.

## 3 RESULTS

Our data curation criteria resulted in 3,481 samples (1,277 controls, 297 influenza infection, 1,907 influenza vaccinated, 1,537 males and 1,944 females) (see SDF1 of online supplementary data files). Our 3,481 samples are from 1,147 individuals. Some studies include repeated measures (in the curated studies individuals were followed for several days after vaccination or infection and varying timepoints were reported as a different samples for the same subject). We included all repeated measures in our downstream analysis and accounted for them in our model. The main results are summarized below, and further discussed in the Discussion Section 4.

**Table 2.** Enriched KEGG Pathways from Statistically Significant Genes with an Interaction Between Disease Status and Age.

| KEGG ID | KEGG Pathway | Gene Count | *p*-value | *adjusted p*-value |
| --- | --- | --- | --- | --- |
| path:hsa04060 | Cytokine-cytokine receptor interaction | 34 | 5.5E-09 | 1.4E-06 |
| path:hsa04660 | T cell receptor signaling pathway | 19 | 6.9E-08 | 8.7E-06 |
| path:hsa04650 | Natural killer cell mediated cytotoxicity | 19 | 3.8E-06 | 3.2E-04 |
| path:hsa04672 | Intestinal immune network for IgA production | 11 | 5.9E-06 | 3.8E-04 |
| path:hsa04640 | Hematopoietic cell lineage | 14 | 1.8E-05 | 9.1E-04 |
| path:hsa05340 | Primary immunodeficiency | 8 | 1.3E-04 | 4.8E-03 |
| path:hsa04064 | NF-kappa B signaling pathway | 13 | 1.4E-04 | 4.8E-03 |
| path:hsa04622 | RIG-I-like receptor signaling pathway | 11 | 1.6E-04 | 4.8E-03 |
| path:hsa04068 | FoxO signaling pathway | 16 | 1.7E-04 | 4.8E-03 |
| path:hsa05166 | HTLV-I infection | 23 | 6.4E-04 | 1.5E-02 |
| path:hsa05162 | Measles | 15 | 6.4E-04 | 1.5E-02 |
| path:hsa04062 | Chemokine signaling pathway | 18 | 9.8E-04 | 2.1E-02 |
| path:hsa05330 | Allograft rejection | 7 | 1.1E-03 | 2.2E-02 |
| path:hsa04380 | Osteoclast differentiation | 14 | 1.4E-03 | 2.6E-02 |
| path:hsa05320 | Autoimmune thyroid disease | 8 | 1.9E-03 | 3.2E-02 |
| path:hsa04110 | Cell cycle | 13 | 2.3E-03 | 3.6E-02 |
| path:hsa04010 | MAPK signaling pathway | 21 | 2.8E-03 | 4.1E-02 |
| path:hsa04630 | Jak-STAT signaling pathway | 15 | 2.9E-03 | 4.1E-02 |
| path:hsa05164 | Influenza A | 16 | 3.3E-03 | 4.4E-02 |

### 3.1 Differentially Expressed Genes in Influenza Disease and Vaccination

Filtering our LRT analysis results by disease factor (see SDF3 of online supplementary data files) for Bonferroni adjusted p-values (*<* 0.05), we identified 4,889 statistically significant disease genes (see SDF5 of online supplementary data files). We performed GO enrichment analysis using BINGO in Cytoscape (version 3.7) (31, 32) and pathway enrichment analysis on the 4,889 genes (Supplementary Figures 1-5 and see SDF6-SDF8 of online supplementary data files). We identified enriched GO terms such as: cell cycle checkpoint (51 genes), response to stimulus (987 genes), immune response (243 genes), transcription (122 genes), regulation of T-cell activation (62 genes), regulation of defense response to virus by host (8 genes) and immune system process (379 genes) (see SDF8 of online supplementary data files for full table). We found 75 enriched KEGG pathways (SDF6 of online supplementary data files). The enriched KEGG pathways include: Cell cycle (68 gene hits), Hematopoietic cell lineage (45 genes), NF-kappa B signaling pathway (46 genes), Metabolic pathways (341 genes), Primary immunodeficiency (23 genes), T cell receptor signaling pathway (44 genes), B cell receptor signaling pathway (29 genes) and also Influenza A (52 genes). We also highlighted the NF-kappa B signaling pathway and the Influenza A KEGG pathways that are relevant to disease with our calculated estimates which compared influenza infection and vaccination expression to that of healthy controls (Figures 4–7).

In addition, we filtered the 4,889 genes for effect size to determine biological significance of the genes (SDF4 of online supplementary data files). We used a two-tailed 10% and 90% quantile filter on the 4,889 genes to: (i)analyze the influenza disease estimates (compared expression to control) list to identify genes that are biologically significant and statistically significant (Bonferroni-adjusted p-value *<*0.05) in influenza infection (ii) analyze the influenza vaccination estimates with the same filtering approach to also identify significant genes for influenza vaccination. For influenza infection our 10% and 90% quantile cut-offs for biological significance were ≤ −0.6724464 and ≥ 0.5949655 respectively. For influenza vaccination, the 10% and 90% quantile cut-offs were ≤ −0.07157763 and ≥ 0.06719048 respectively. For influenza infection we identified 978 genes of the 4,889 to be biologically significant (Table 3 and SDF9 of online supplementary data files), and we also identified 978 genes to be biologically significant for influenza vaccination (Table 3 and SDF10 of online supplementary data files). We then compared the two gene lists to identify the intersection (genes in common), genes only in the influenza disease list, and genes only in the influenza vaccination list (Figure 1D and SDF11-SDF13 of online supplementary data files). There were 334 genes in common across both lists (influenza disease and vaccination) (SDF17 of online supplementary data files) that resulted in enriched Reactome pathways such as Interferon alpha/beta signaling (14 genes), Interferon gamma signaling (12 genes), Antiviral mechanism by IFN-stimulated genes (9 genes), and Cell Cycle Checkpoints (17 genes) (SDF20 of online supplementary data files). There were 644 genes that were only in influenza infection list (SDF18 of online supplementary data files) that resulted in enriched Reactome pathways including: Neutrophil degranulation (45 genes), Cell Cycle Checkpoints (27 genes), Amplification of signal from the kinetochores (13 genes), Amplification of signal from unattached kinetochores via a MAD2 inhibitory signal (13 genes) and Mitotic Spindle Checkpoint (14 genes) (SDF21 of online supplementary data files). Also, we identified another 644 genes that were only in the biologically significant list for the vaccinated subjects (SDF19 of online supplementary data files). Enriched Reactome pathway analysis on these genes resulted in pathways such as Interferon Signaling (24 genes), Antigen processing-Cross presentation (14 genes), ER-Phagosome pathway (12 genes), Binding and Uptake of Ligands by Scavenger Receptors (8 genes) and Class I MHC mediated antigen processing & presentation (30 gene) (SDF22 of online supplementary data files).

**Table 3.** Top 10 Up- and Down-Regulated Differentially Expressed Genes from the Influenza Infected and Influenza Vaccination Biologically Significant Gene Lists (based on estimates).

| Influenza Infection |  |  |  |
| --- | --- | --- | --- |
| Down-Regulated |  | Up-Regulated |  |
| Gene | Difference of Means | Gene | Difference of Means |
| NELL2 | -1.687 | UGCG | 2.005 |
| UBASH3A | -1.583 | CD177 | 1.875 |
| ABCB1 | -1.513 | OTOF | 1.844 |
| PID1 | -1.457 | HP | 1.625 |
| CACNA2D3 | -1.428 | SSH1 | 1.491 |
| PTGDR | -1.423 | DTL | 1.431 |
| CD40LG | -1.392 | GPR84 | 1.428 |
| PTGDR2 | -1.390 | HJURP | 1.420 |
| TLE2 | -1.379 | CDC45 | 1.395 |
| NCR3 | -1.357 | SLC1A3 | 1.390 |
| Influenza Vaccination |  |  |  |
| Down-Regulated |  | Up-Regulated |  |
| Gene | Difference of Means | Gene | Difference of Means |
| TOP1MT | -0.179 | GBP1 | 0.354 |
| ARNTL | -0.176 | MYOF | 0.347 |
| DIDO1 | -0.172 | STAT1 | 0.284 |
| PDE4D | -0.169 | PSTPIP2 | 0.281 |
| TMX4 | -0.168 | SAMD9L | 0.276 |
| ZNF589 | -0.166 | OAS3 | 0.269 |
| SLC37A3 | -0.165 | WARS | 0.263 |
| GNB5 | -0.165 | BATF2 | 0.263 |
| ENO2 | -0.162 | ANKRD22 | 0.256 |
| AP3M2 | -0.159 | C1QB | 0.255 |

We also explored the 4,889 genes to identify how many genes were different in gene expression when looking at influenza infected subjects compared to influenza vaccinated subjects. Of the 4,889 genes, 4,261 genes showed statistically significant differences between vaccination and infection with influenza (Figure 3 and SDF25 - SDF27 of online supplementary data files).

**Figure 3.**
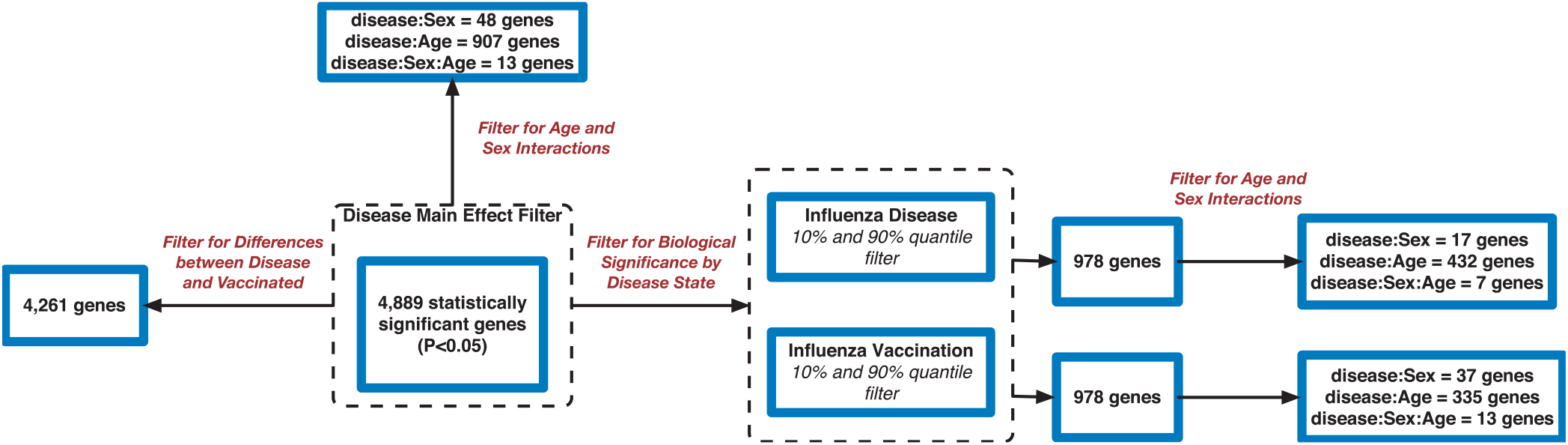
Flowchart of Gene Filtering Steps for Influenza Meta-analysis.

### 3.2 Age and Sex Effect on Gene Expression in Influenza

Using the 4,889 genes disease significant genes from above, we Bonferroni-adjusted the p-values for both the age and sex factors. We then further filtered the 4,889 list by the age factor p-values (Bonferroni-adjusted p-value *<* 0.05) to identify statistically significant interacting genes between disease state and age (DxA). We also repeated this approach for the sex factor interaction with disease (DxS). Of the 4,889 statistically significant (Bonferroni-adjusted p-value *<*0.05) disease genes, 907 of them had a statistically significant interaction with disease and age (SDF28 of online supplementary data files). KEGG enrichment, our results include: Cytokine-cytokine receptor interaction (34 genes), T cell receptor signaling pathway (19 genes), Natural killer cell mediated cytotoxicity (19 genes), Intestinal immune network for IgA production (11 gene hits), Hematopoietic cell lineage (14 genes), Primary immunodeficiency (8 genes), NF-kappa B signaling pathway (13 genes), and Influenza A (16 genes) (SDF30 of online supplementary data files). We also looked at the biologically significant gene lists for influenza infection and vaccination (based on effect as discussed above) to determine which of these genes also had a significant interaction with disease and age. Of the 978 in the influenza infection biologically significant list, 432 had a statistically significant ((Bonferroni-adjusted p-value *<* 0.05 for disease and age factor) interaction with disease and age (Figure 3 and SDF32 of online supplementary data files). In the biologically significant gene list for influenza vaccinated subjects 335 genes also had a statistically significant (Bonferroni-adjusted p-value *<* 0.05 for disease and age factor) interaction with disease and age (Figure 3 and SDF35 of online supplementary data files).

Furthermore, we explored differences in gene expression (based on mean differences across groups) in subjects with influenza infection, influenza vaccination and controls across the 4 age groups: (−1,3],(3,19], (19,65], (65,100] using the gene lists of identified disease:age interacting genes. First we calculated the mean expression for control subjects younger than 3 (age group: (−1,3]). This served as our baseline for all comparisons to influenza infection and vaccination. We calculated the difference in means for the subjects within the other age groups only focusing on the healthy subjects and used the younger than 3 as our baseline to find the difference in means (Figure 8). We also calculated the difference in mean expression for all influenza infected subjects and used the influenza infected subjects younger than 3 as the baseline for comparisons of relative expression (Figure 9A). In addition to this, we calculated difference in means by comparing influenza infected samples to the control baseline (younger than 3) (Figure 9B). We repeated the above steps with our vaccinated subjects to explore how expression changes with age and disease state (Figure 10). We also plotted the difference in means comparing influenza vaccinated subjects to influenza infected subjects to highlight temporal patterns of the 907 interacting (disease:age) genes (Figure 11).

We also filtered our gene lists (statistically significant disease genes and the biologically significant for influenza disease and vaccination gene lists) for genes with a statistically significant disease interaction with sex (Figure 3). We identified 48 of the 4,889 disease genes (Bonferroni-adjusted p-value *<* 0.05 for disease and sex factor) that interacted with disease and sex ((Figure 3) and SDF29 of online supplementary data files). In the influenza infected biologically significant gene list there were 17 genes that interacted with disease and sex (Bonferroni-adjusted p-value *<* 0.05 for disease and sex factor), and 7 genes had an interaction with disease, sex and age (Bonferroni-adjusted p-value *<* 0.05 for disease, sex and age factor) (Figure 3 and see also SDF33 and SDF34 of online supplementary data files). We did not find any statistically significant enrichment in pathways for these genes. As for the biologically significant influenza vaccination genes, 37 of them were associated with disease and sex interactions (Bonferroni-adjusted p-value *<* 0.05 for disease and sex factor), and 13 genes had associated interactions with disease, sex and age (Bonferroni-adjusted p-value *<* 0.05 for disease, sex and age factor) (Figure 3 and see also SDF36 and SDF37 of online supplementary data files). We also did not find any enriched pathways for these genes.

## 4 DISCUSSION

Every year there is a new vaccine available to reduce the amount of influenza cases worldwide. The influenza virus is constantly changing and researchers have to predict the most common strains that will affect the population each season. During the flu season, the majority of hospitalizations and deaths from influenza are within the elderly population (3). Young children are also at high risk for severe infections of influenza due to their underdeveloped immune system (33). Current vaccine development methods, though effective are also flawed. In some cases, the influenza strains can mutate after the strains for the vaccine have been selected for the upcoming flu season, which then reduces the effectiveness of the vaccine (34). Exploring how gene expression varies in influenza infection, vaccination, and comparison of the differences may highlight prospective biomarkers/gene signatures for improving vaccinations. In addition, because of the observed age-dependency in influenza infection, investigating gene expression temporal patterns across various ages can also provide insight on how genes change due to underdevelopment and immunosenescence.

We identified 18 microarray expression datasets that passed our inclusion criteria for a meta-analysis on influenza (Table 1). We collected the raw expression microarray data for all datasets, pre-processed them and combined by common gene names. With 3,481 samples (including repeated measures) we modeled the pre-processed expression data with a mixed effects model and carried out LRT analysis. Our LRT analyses resulted in 4,889 statistically significant (Bonferroni-adjusted p-value *<*0.05) disease genes (see SDF5 of online supplementary data files). These results include CD177 which plays a role in innate immune response by regulating chemotaxis of neutrophils (35, 36), BCL11B which regulates T-cell differentiation (37, 36), HMGB1 protein has been shown to promote viral replication (38) and plays a role in inflammation (36), TPP2 plays a role in major histocompatibility complex (MHC) presentation and TANK is involved in NF-kappa B signalling.

We highlighted the KEGG NF-kappa B signaling pathway using the estimates from influenza infection and vaccination (Figure 4) and Figure 5). The NF-kappa B pathway is activated during influenza infection which up-regulates antiviral genes (39) and can regulate viral synthesis (40). Previous studies have also reported that the influenza virus is capable of regulating antiviral activity by NF-kappa B and promote replication in hosts (40). In the NF-kappa B pathway, we observed similar expression patterns between disease and vaccinated subjects, including down regulation of genes involved in MCH/Antigen presentation for both physiological states. There are also some differences in gene expression observed such as CD40 and PARP1 up-regulated in vaccinated samples. CD40 has previously been shown to regulate immune response and promotes protection against the virus (41, 42) while PARP1 has been highlighted as a host factor that can regulate the polymerase activity of influenza (43). In Figure 5, the genes in our vaccine list in the RIG-I-like receptor signaling pathway are down-regulated, compared to influenza infected subjects (Figure 4). The RIG-I-like receptors have been previously shown to be involved in sensing viral RNA and regulating an antiviral immune response (44). Other genes such as ICAM which is involved in lymphocyte adhesion and T-cell costimulation, and BLC and ELC involved in lymphoid tissue honing are all down-regulated in vaccinated subjects compared to infected subjects (Figure 4) and Figure 5). We also highlighted expression of genes in the Influenza A KEGG pathway for influenza infected and influenza vaccinated (Figure 6) and Figure 7). Although there are similarities in both figures, some key differences in expression are observed in genes connected with high fever (IL-1 and IL-6). Studies have shown elevated levels of IL-1β and IL-6 following infection with influenza A (45, 46).

**Figure 4.**
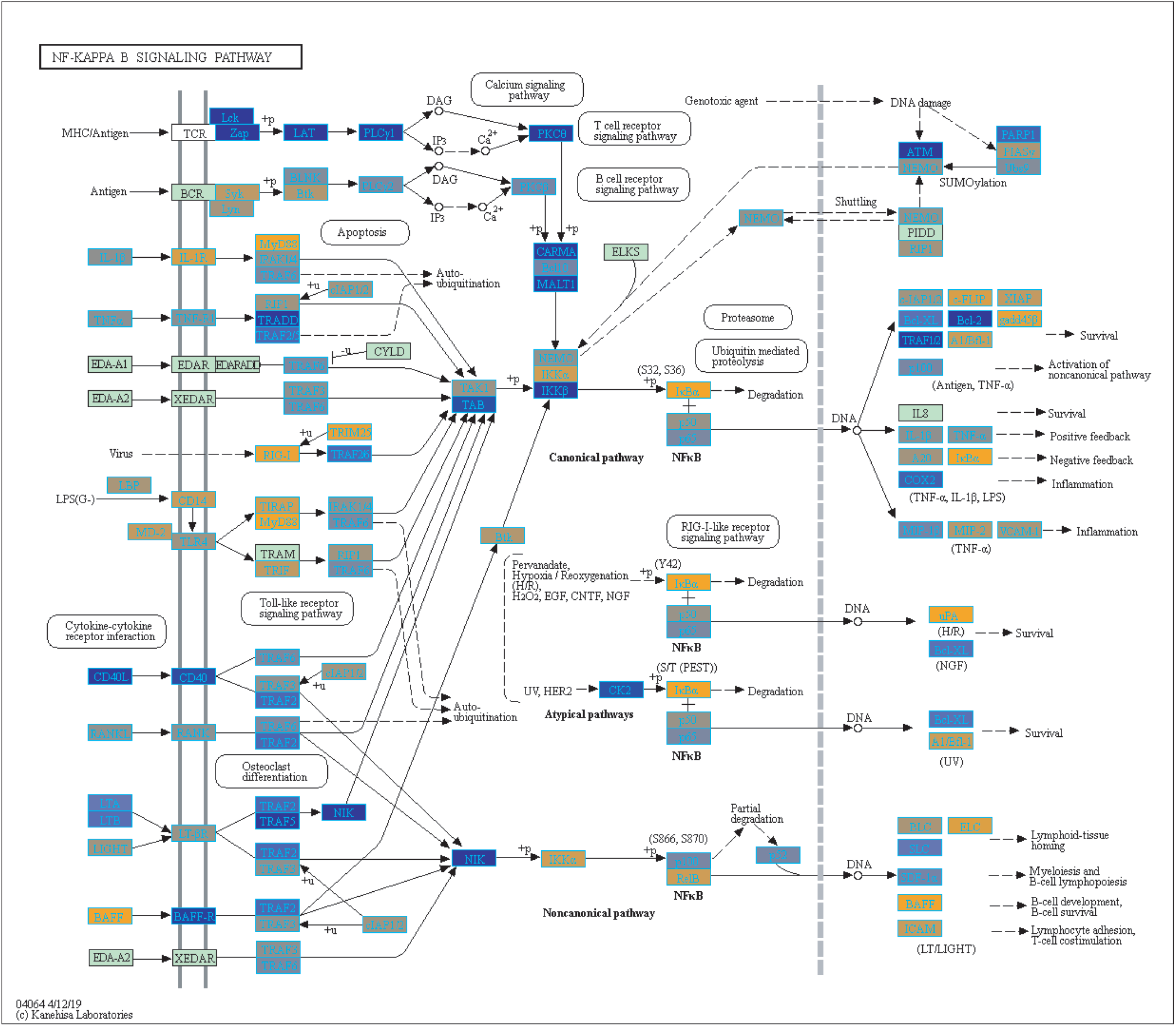
Highlighted NF-Kappa B Signaling KEGG Pathway (hsa04040) with Enriched Genes from the LRT Analysis (Bonferroni-adjusted p-value *<* 0.05) for Influenza Infected Subjects (67, 68, 69). Yellow-colored genes are up-regulated and blue-colored genes are down-regulated in Influenza Infected Subjects.

**Figure 5.**
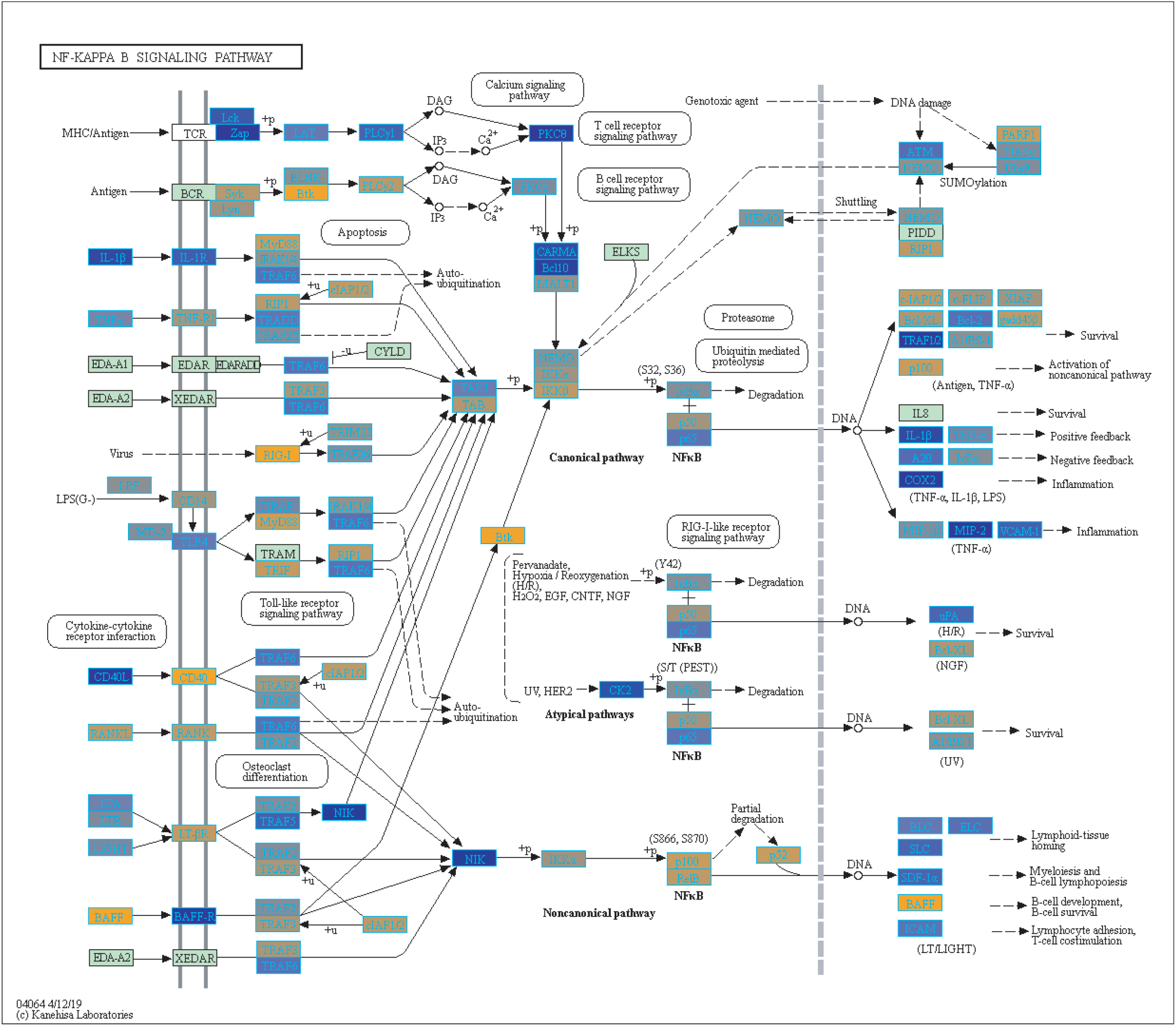
Highlighted NF-Kappa B Signaling KEGG Pathway (hsa04040) with Enriched Genes from the LRT Analysis (Bonferroni-adjusted p-value *<* 0.05) for Influenza Vaccinated Subjects (67, 68, 69). Yellow-colored genes are up-regulated and blue-colored genes are down-regulated in Influenza Vaccinated Subjects.

**Figure 6.**
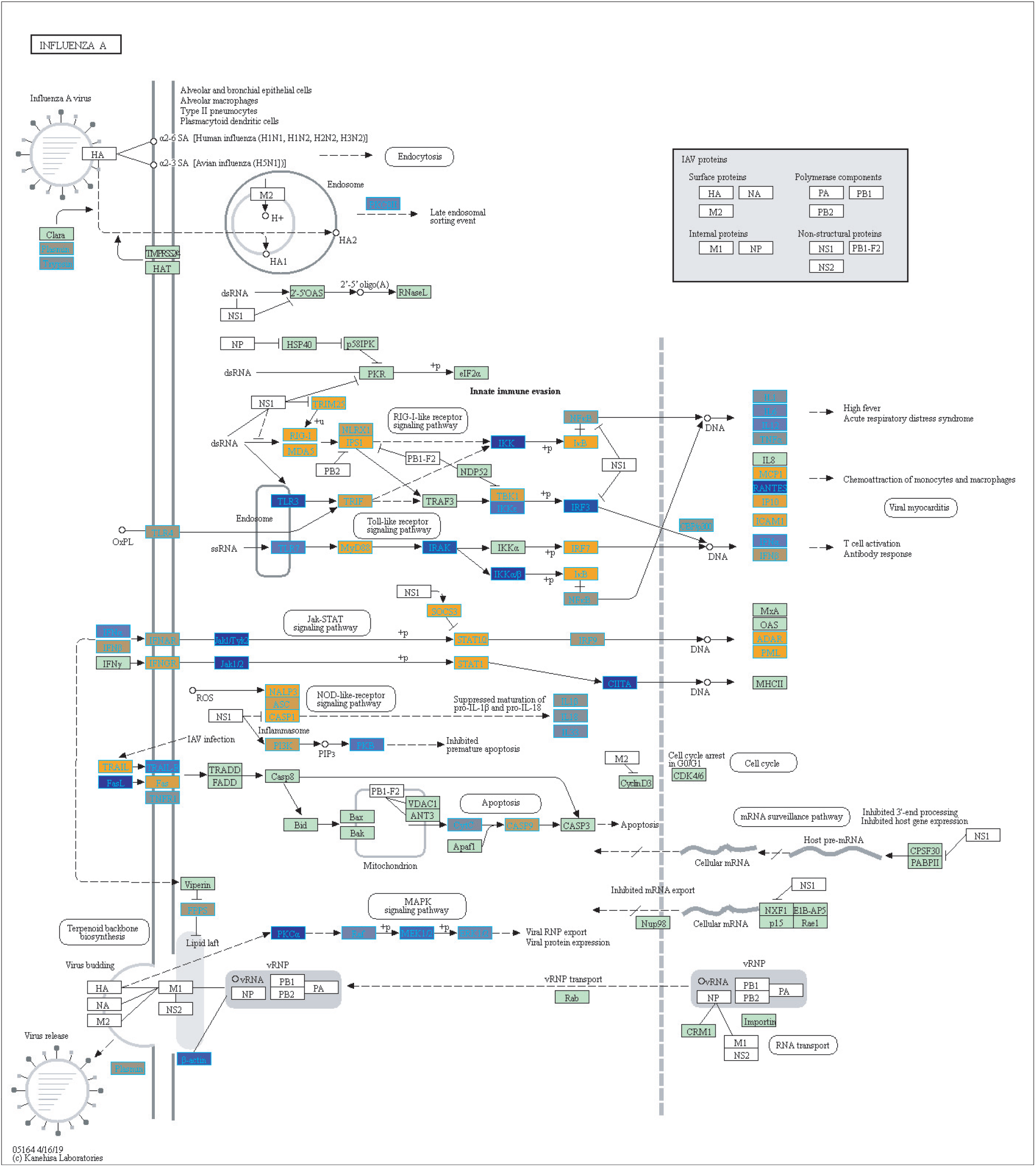
Highlighted Influenza A KEGG Pathway (hsa05164) with Enriched Genes from the LRT analysis (Bonferroni-adjusted p-value *<* 0.05) for Influenza Infected Subjects (67, 68, 69). Yellow-colored genes are up-regulated and blue-colored genes are down-regulated in Influenza Infected Subjects.

**Figure 7.**
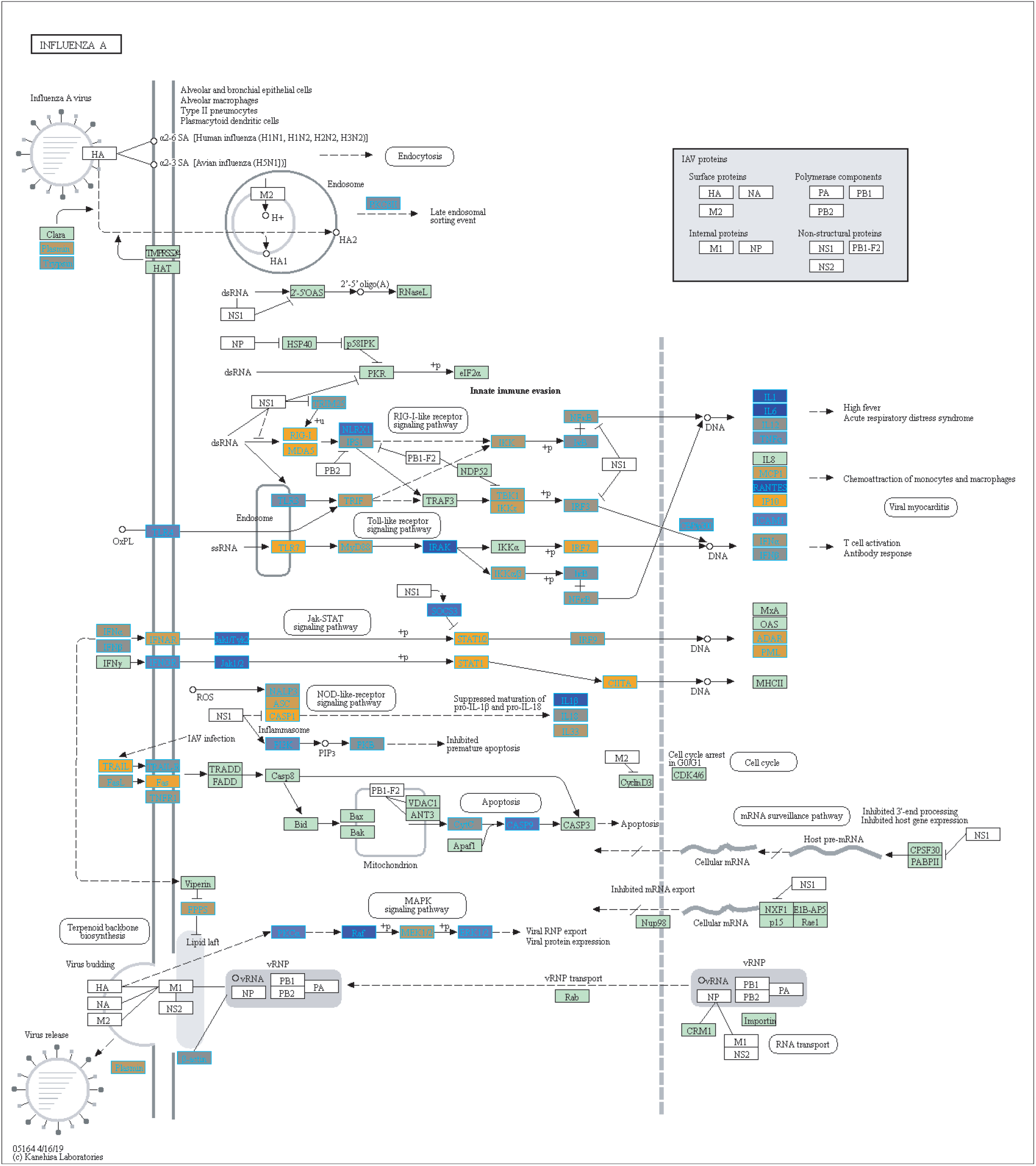
Highlighted Influenza A KEGG Pathway (hsa05164) with Enriched Genes from the LRT Analysis (Bonferroni-adjusted p-value *<* 0.05) for Influenza Vaccinated Subjects (67, 68, 69). Yellow-colored genes are up-regulated and blue-colored genes are down-regulated in Influenza Vaccinated Subjects.

Additionally, we compared our biologically significant gene list for influenza infection (978 genes) to Dunning et al.,who identified whole blood RNA signatures in hospitalized adults with influenza. Their findings indicated that genes involved in interferon-related pathways were activated at the start of the infection and by day 4 had started to decrease with a shift in inflammatory and neutrophil related pathways (46). Our findings also indicate enrichment for neutrophil related pathways in the case of influenza infection. Dunning et al. list a top 25 gene set (controls versus influenza subjects), from which 22 genes overlap with our findings (978 genes, see SDF9 of online supplementary data files). 5 of our top 10 up-regulated gene list overlap with the Dunning et al. 25-gene set (Table 3), namely UGCG, CD177, OTOF, HP and SSH1.

Furthermore, our identified biologically significant gene lists for influenza infection and vaccination (using a 2-tailed 10% quantile filter on expression estimates of effect size compared to healthy control) have 334 genes in common, with 644 genes being unique to influenza infection and 644 being unique to influenza vaccination (SDF17-SDF19 of online supplementary data files). Following pathway enrichment, we observed that the genes that are unique to each disease state (influenza infected and vaccinated) are involved in different processes. For example, the biologically significant genes only in influenza infected samples were enriched in pathways such as neutrophil degranulation and cell cycle checkpoints (SDF21 of online supplementary data files). Neutrophil degranulation is a defensive process neutrophils undergo to protect the host against invading pathogens. On the other hand, pathways involved in interferon signaling and antigen processing were enriched for the genes only in the vaccinated gene list (SDF22 of online supplementary data files). This indicates that with the actual infection the body undergoes different processes to that induced by vaccination (Supplementary Figures 3-4 and see SDF23 and SDF24 of online supplementary data files).

The 48 genes for which we identified a statistically significant interaction between disease and sex are highlighted in SDF29 of the online supplementary data files. Sex-specific gene expression has been previously observed in influenza. Studies have observed females exhibited a stronger immune response to influenza vaccine compared to males within the first day (47). Another study suggested that males have a stronger immune response to influenza infection (48). These findings indicate sex and influenza is still to be explored and our gene list may offer new candidates to be investigated for their role in influenza.

With regards to aging and influenza, the statistics of the disease burden indicates specific age groups are at higher risk for infection (3). This is in part due to immune system development and deterioration. For example, B and T cell function diminishes with age (49, 33). In our analysis, we identified 907 disease-associated genes with a statistically significant interaction with age that were also enriched in immune related KEGG pathways (SDF30 of online supplementary data files). Figure 8 compares the mean differences of healthy subjects to the baseline (healthy children younger than 3). There are 4 major groups (Figure 8 and see SDF47 of online supplementary data files): with reference to Figure 8, genes in Cluster 1 were up-regulated compared to the baseline for all age comparisons, Cluster 2 and 3 genes were generally down-regulated compared to the baseline, and Cluster 4 genes are up-regulated and increase with age. Genes in Cluster 1 and 2 are involved in Reactome pathways such as cytokine signaling, interferon signaling and the immune system. Cluster 3 genes are involved in Reactome pathways such as interferon signaling and cell cycle while Cluster 4 genes are involved in cellular senescence, signaling by interleukins and immune system.

**Figure 8.**
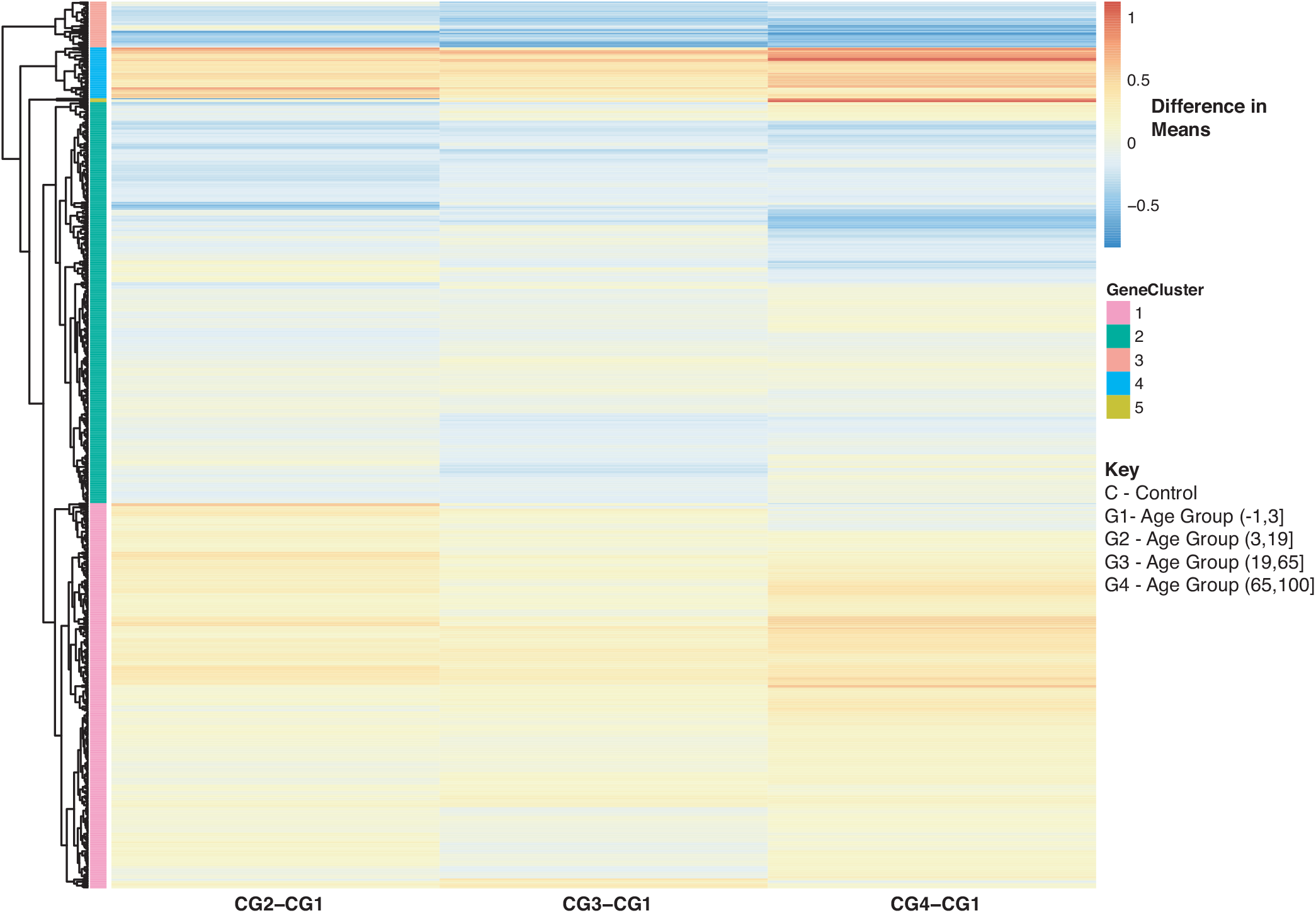
Heatmap of Statistically Significant (Bonferroni-adjusted p-value *<*0.05) Genes with an Interaction Between Disease State and Age for Healthy Controls. Difference in means calculated by comparing control subjects in age groups 2-4 to control subjects in age group 1 (baseline).

In Figure 9 we further explored changes in gene expression across age groups due to influenza infection of our 907 disease:age interacting genes. Figure 9A is compares influenza infected subjects in age groups 2,3 and 4 to the baseline (infection subjects under 3). In Figure 9A there are three major groups (cluster numbering with respect to Figure 9A): Cluster 1 (gradual decrease with age), 2 (genes up-regulated with increase in age), and 3 (gradual down-regulation with age). Genes in Cluster 1 are in Reactome pathways such as cytokine signaling, interferon signaling, antiviral mechanism by IFN-stimulated genes and chemokine receptors. Cluster 2 genes are involved in regulatory T lymphocytes, transcription, protein repair, and interleukin-2 signaling while Cluster 3 genes are involved in gene transcription (see SDF48 of online supplementary data files). Figure 9B instead compares influenza infected subjects to controls by looking at difference in means. There are three groups of expression patterns (cluster numbering with respect to Figure 9B): Cluster 1 shows a gradual increase with age, in Cluster 2 expression intensifies with age and in Cluster 3 genes are down-regulated compared to the control subjects younger than 3 (see SDF49 of online supplementary data files). Genes in Clusters 1 and 2 were not in any enriched Reactome pathways but are associated with transcription and signaling pathways. Genes in Cluster 3 were in Reactome pathways that include cytokine signaling, interferon signaling, antiviral mechanism by IFN-stimulated genes and chemokine receptors.

**Figure 9.**
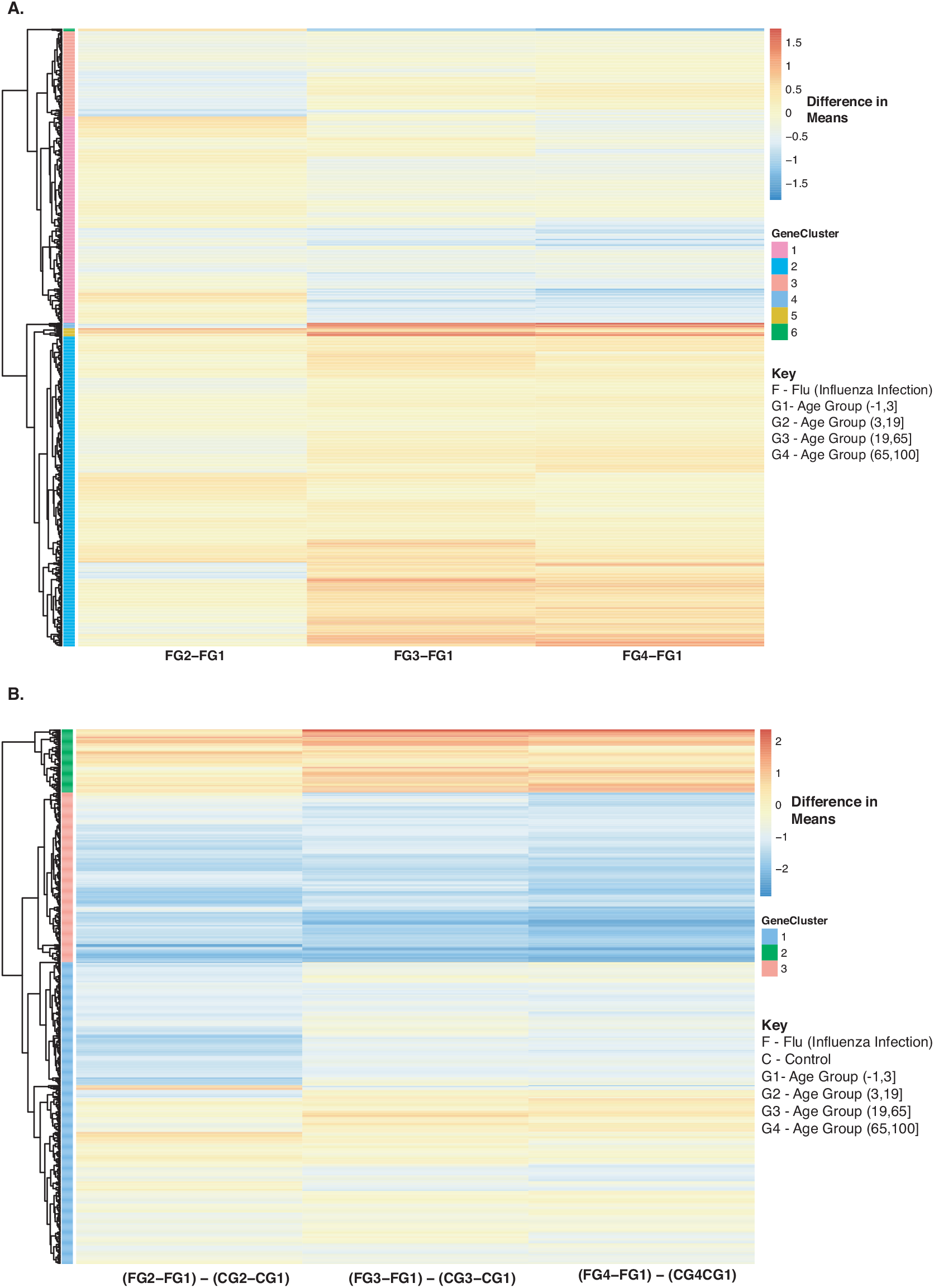
Heatmap of Statistically Significant (Bonferroni-adjusted p-value *<*0.05) Genes with an Interaction Between Disease State and Age for Influenza Infected Subjects. (A)Difference in means calculated by comparing influenza infected subjects in age groups 2-4 to influenza infected subjects in age group 1 (baseline). (B) Comparison of influenza infected subjects to control subjects in the different age groups by calculating the difference between the baseline-adjusted means for influenza infected subjects (A) and control subjects (Figure 8).

As for the vaccinated subjects with respect to Figure 10A, we observe a gradual decrease in gene expression for gene Cluster 2 and a gradual increase in expression for genes in Cluster 1 and 3 compared to the baseline (young vaccinated subjects under age 3) (SDF50 of online supplementary data files). Genes in Cluster 1 were not enriched in pathways while genes in Cluster 2 were enriched in Reactome pathways that include interferon and cytokine signaling, antiviral mechanism and response. Genes in Cluster 3 were enriched in Reactome pathways that include interferon and cytokine signaling, cellular senescence and immune system. When we compared vaccinated subjects to control subjects across ages we observed 3 main trends, with respect to Figure 10B: Cluster 1 (pathways include antiviral mechanisms, interferon and cytokine signaling) 2 (pathways such as immune response and cell migration, immunological synapse and chemokine receptors) and 4 (mitochondrial translation) genes are all up-regulated in vaccinated subjects with Cluster 3 (pathways include interferon and cytokine signaling and immune system) genes being down-regulated (Figure 10B and SDF51 of online supplementary data files).

**Figure 10.**
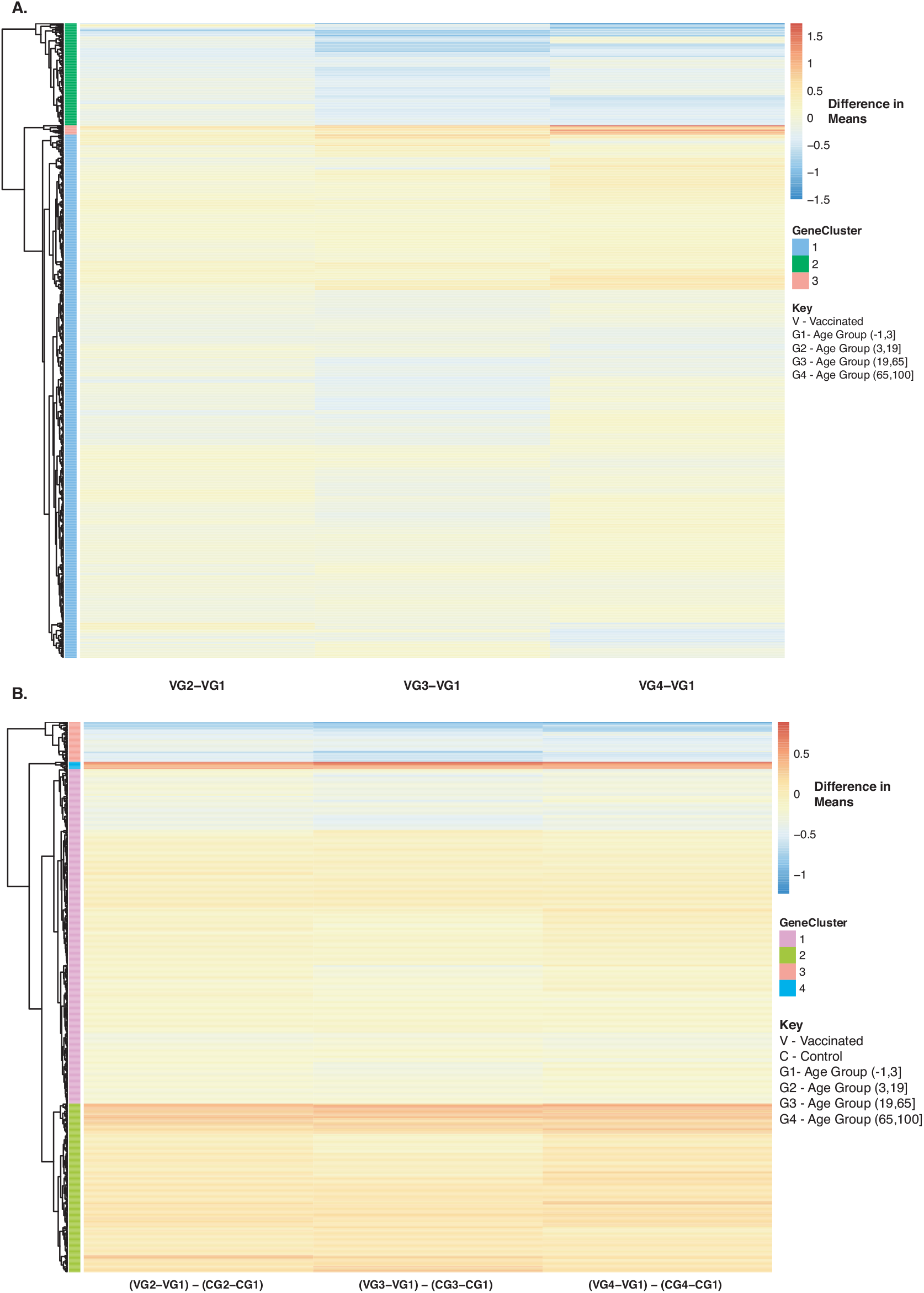
Heatmap of Statistically Significant (Bonferroni-adjusted p-value *<*0.05) Genes with an Interaction Between Disease State and Age for Influenza Vaccinated Subjects. (A)Difference in means calculated by comparing influenza vaccinated subjects in age groups 2-4 to influenza vaccinated subjects in age group 1 (baseline). (B) Comparison of influenza vaccinated subjects to control subjects in the different age groups by calculating the difference between the baseline-adjusted means for influenza vaccinated subjects (A) and control subjects (Figure 8).

**Figure 11.**
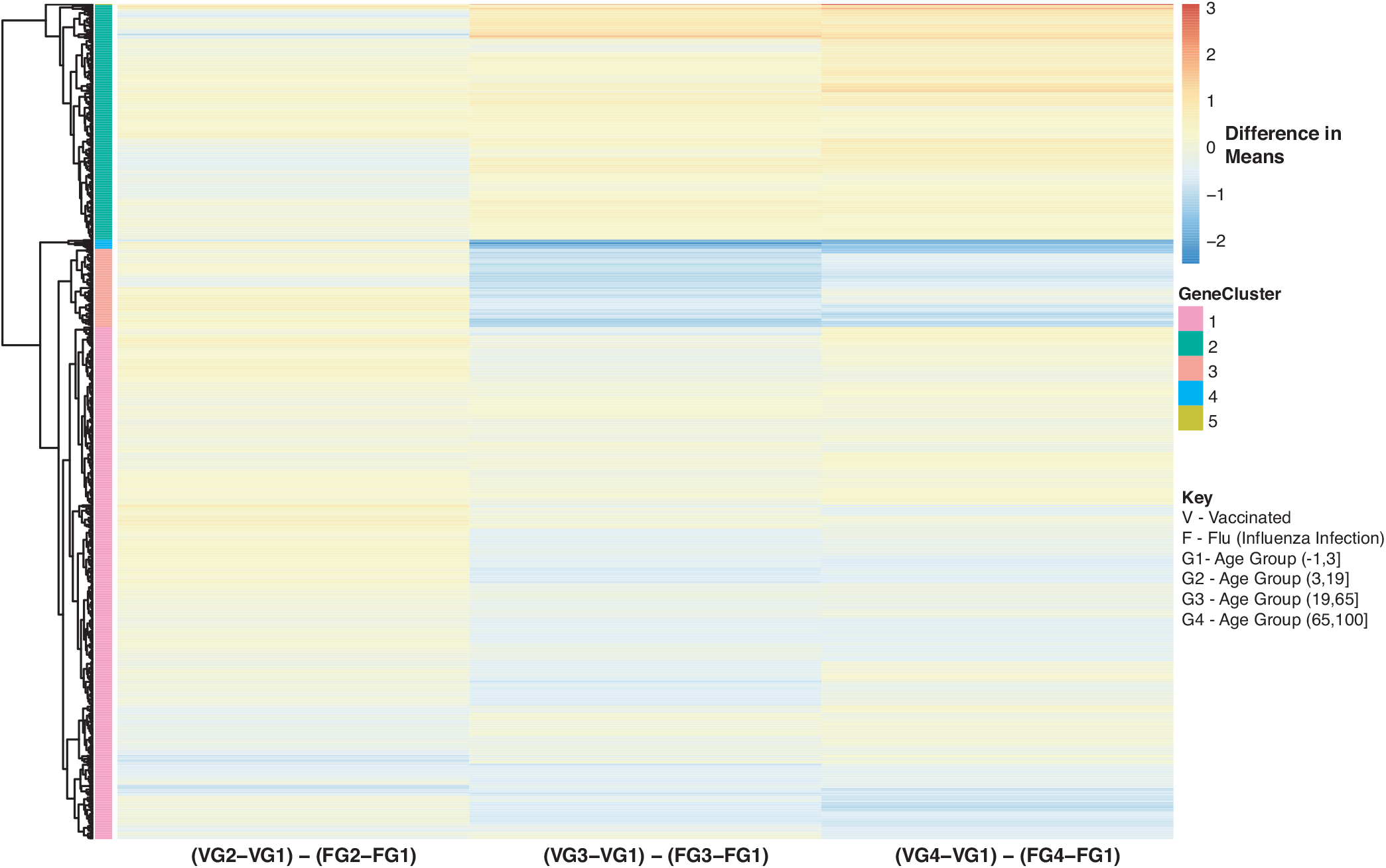
Heatmap of Statistically Significant (Bonferroni-adjusted p-value *<*0.05) Genes with an Interaction Between Disease State and Age for Influenza Vaccinated Subjects Compared to Influenza Infected Subjects. Comparison of baseline-adjusted means for influenza vaccinated subjects (Figure 10A) and influenza infected subjects (Figure 9A).

We also compared influenza vaccinated subjects to influenza infected subjects to explore changes in expression with age. There are 3 major groups (cluster numbering with respect to Figure: Cluster 1 and 3 genes show a gradual decrease in expression with age in vaccinated subjects compared to influenza infected subjects. Genes in Cluster 2 show a gradual increase in expression with age in influenza vaccinated subjects. Supplementary Figures 6-8 also explore temporal patterns with age.

Our heatmaps display temporal patterns with age in response to influenza infection and vaccination. These genes that are associated with disease and age interactions, are all involved in immune-related pathways. Exploring how gene expression changes with age in immune related genes can help further characterize the disease and improve treatments. For example, age, preexisting health conditions and influenza history (previous infection or vaccination) are all factors that can affect the efficacy of the vaccine (50). There is an on-going effort to improve efficacy of vaccination in the elderly population. Studies have suggested that antibody titers decline drastically in older adults from seroconversion to day 180 after vaccination (50). The decay of antibody titers also highlight the importance of determining the right time and how many times one should be vaccinated. Vaccines for the elderly population have been modified to increase the dosage and use adjuvants to increase immunogenicity (51). and Ramsay et al., also showed that vaccination during the current influenza season provides stronger protection than vaccinations from previous seasons (52).

The vaccine type also plays a role in immunogenicity within hosts. For example, Nakaya et al., were able to detect larger antibody titers and plasmblasts generated in the trivalent inactivated vaccine (TIV) compared to the live attenuated vaccine (LAIV), and differentially expressed genes mostly related to interferon signaling (53). LAIV responses in young children are higher than in adults. For instance, LAIV when compared to inactivated vaccines induced smaller concentrations of antibodies in response to HA in adults (54). Previous findings have shown the benefit of taking a systems biology approach to assess gene expression responses to vaccinations (55, 56, 50). Our findings not only identify genes that are different between controls compared to infected and vaccinated subjects, but with our methodology we were also able to assess differences between the influenza infected and vaccinated subjects while still investigating disease genes that interact with age and sex. Our temporal patterns with age for each disease state helps to clarify how age might be playing a role.

As we have previously observed (16), meta-analyses using microarray expression data have multiple limitations: Our findings are limited to only genes that have been annotated and are existing probes on the arrays, and also have to be consistently utilized across array platforms. Hence, we are unable to probe global gene expression, and are limited to mRNA profiling. These can be expanded in future studies using RNA-sequencing data, and the newer single-cell sequencing approaches that would allow cell-specific information to be discerned, which is important in evaluating immune responses and the interplay between various cell types. Taking a similar approach to our microarray dataset analysis using RNA-sequencing data will promote the discovery of novel genes by being able to explore the entire transcriptome. Additionally, we are limited by the varying annotations of the available public datasets, and can only explore characteristics that are uniformly reported. For example, we did not have virus strain information for all samples or vaccine details so we were unable to include such info in our analysis. In addition to this, our study is unbalanced (particularly with respect to disease state, where a limited number of influenza infection samples were available: 3,481 samples (1,277 controls, 297 influenza infection, 1,907 influenza vaccinated, 1,537 males and 1,944 female). We additionally used repeated measures, which we accounted for in our mixed effects model.

Despite the limitations introduced by using micro-array data, our study identified gene candidates by factor (disease status, age and sex) that can be examined further to understand their role in influenza infection and vaccination. We also highlighted 907 genes that have an age-effect on gene expression. These genes can be further explored to determine their role in influenza infection and how they can be further analyzed for their role in implementing effective universal vaccines regardless of age. All these consideration are of paramount importance in designing the next generation of vaccines, as we move forward towards a universal influenza vaccine.

## AUTHOR CONTRIBUTIONS

All authors listed have made a substantial, direct and intellectual contribution to the work, and approved it for publication.

## SUPPLEMENTAL DATA

Our supplemental figures and tables are provided in the accompanying supplementary data pdf file. All of our datasets/results from our meta-analysis pipeline have been uploaded to FigShare as online supplemental data, as described below (the corresponding online file names begin with a prefix ‘SDF’ and are enumerated as also referred to in the manuscript).

## DATA AVAILABILITY STATEMENT

The datasets generated and analyzed for this study can be found in Figshare’s online digital repository at https://doi.org/10.6084/m9.figshare.8636498.

