## Supplementary Figures for "Microarray Gene Expression Dataset Re-Analysis Reveals Variability in Influenza Infection and Vaccination"

### Supplementary Material

#### 1 SUPPLEMENTARY FIGURES

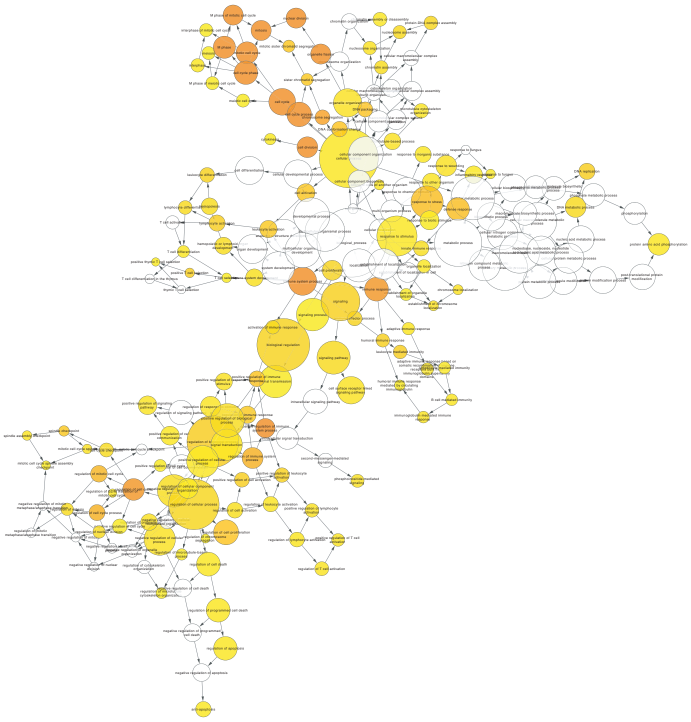

**Figure S1.** Gene Ontology of Biologically Significant Genes for Influenza Infected Subjects using BiNGO. The node size relates to number of genes, and the yellow nodes are statistically significant with a p-value $< 0.05$ and false discovery rate $< 0.05$.

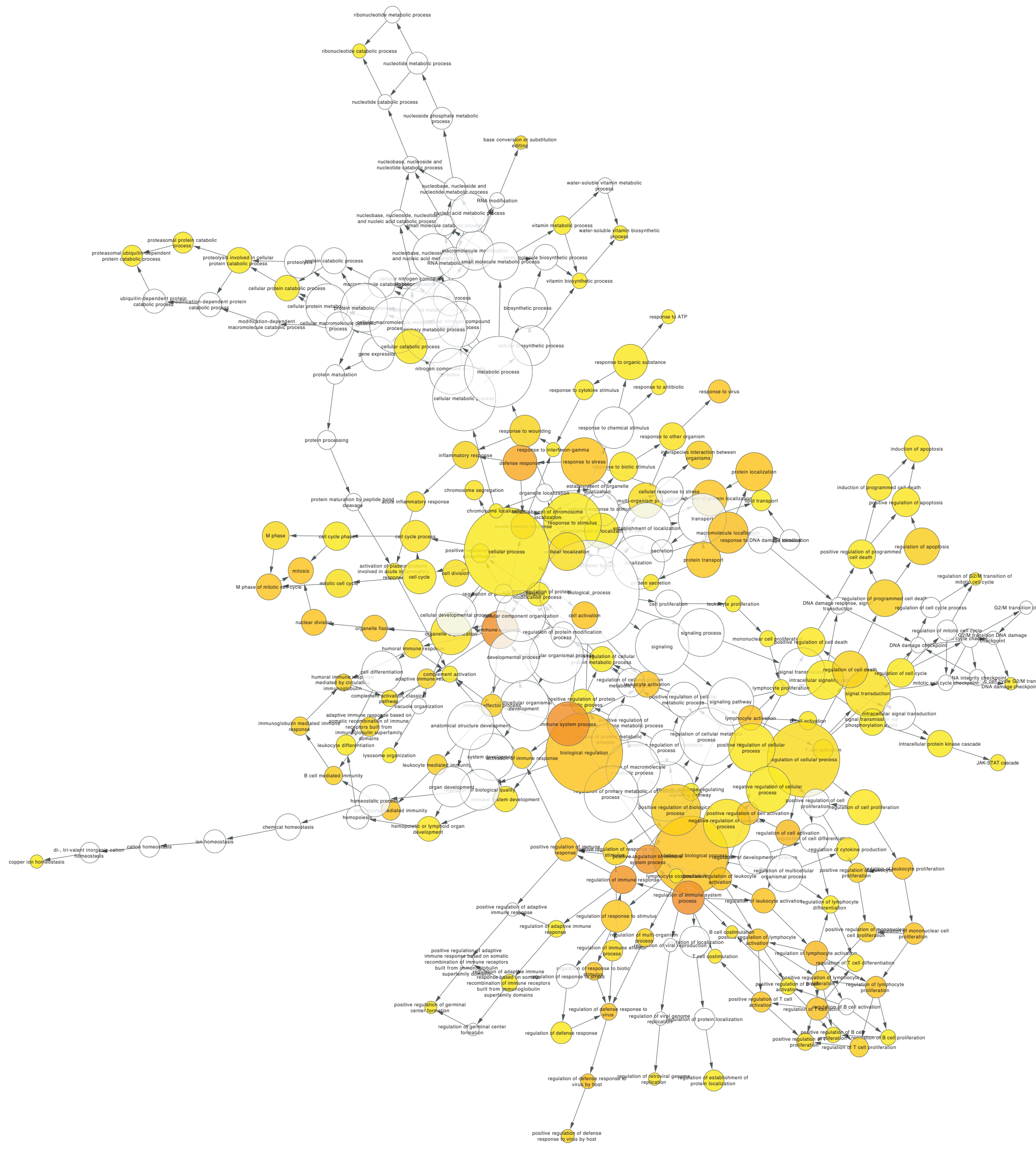

**Figure S2.** Gene Ontology of Biologically Significant Genes for Influenza Vaccinated Subjects using BiNGO. The node size relates to number of genes, and the yellow nodes are statistically significant with a p-value $< 0.05$ and false discovery rate $< 0.05$.

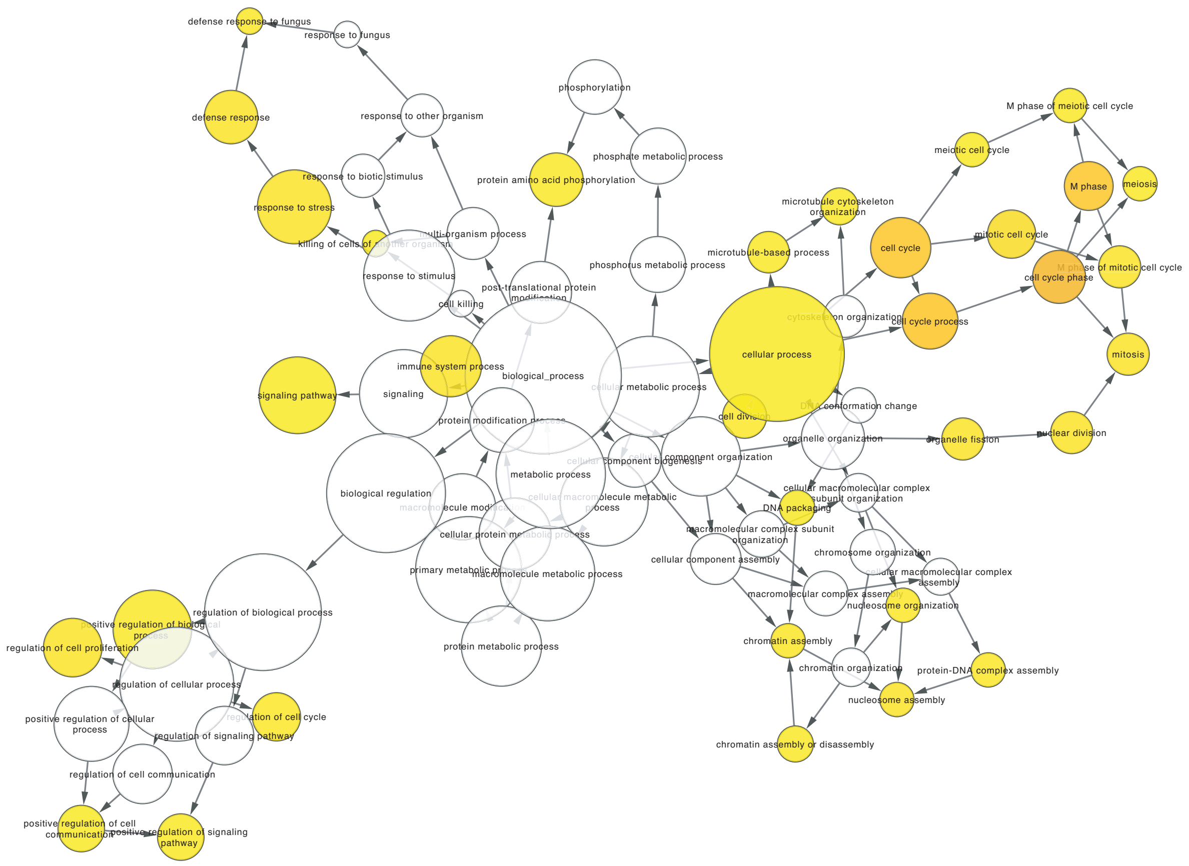


**Figure S3.** Gene Ontology of Biologically Significant Genes Only in the Influenza Infected Subjects Gene List using BiNGO. The node size relates to number of genes, and the yellow nodes are statistically significant with a p-value $< 0.05$ and false discovery rate $< 0.05$.

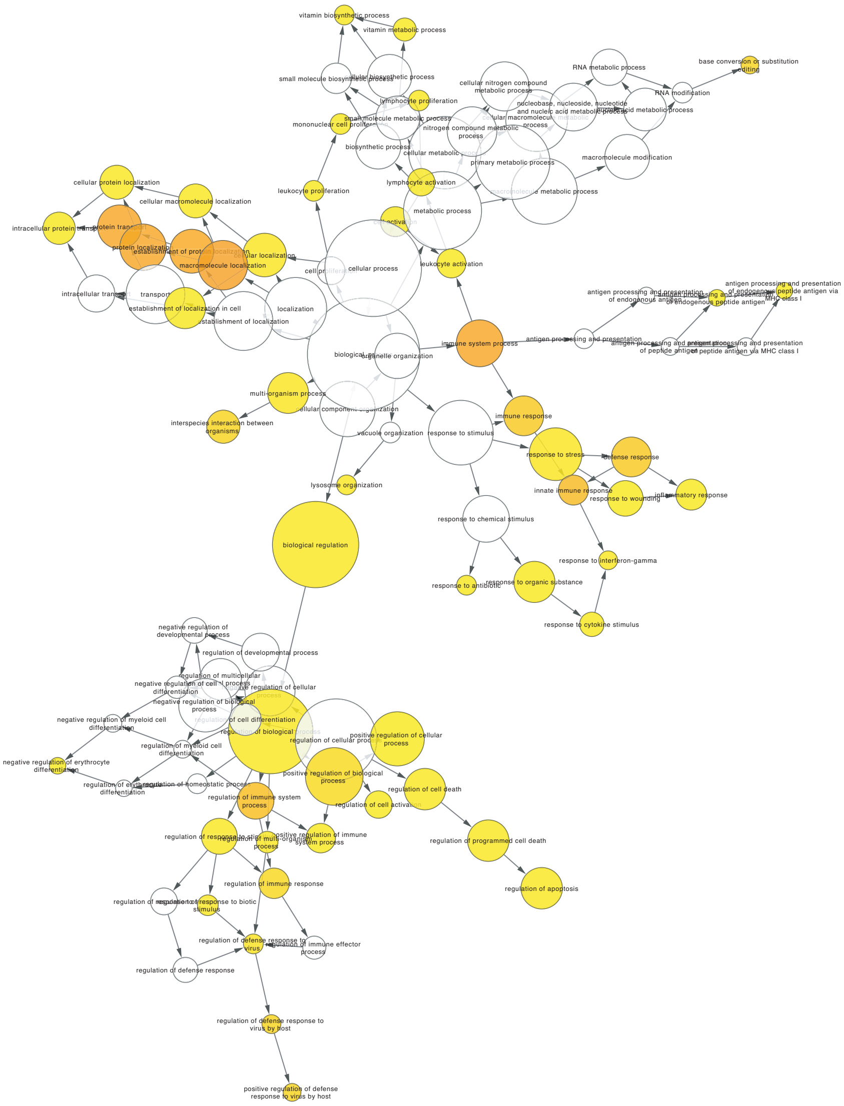


**Figure S4.** Gene Ontology of Biologically Significant Genes Only in the Influenza Vaccinated Subjects Gene List using BiNGO. The node size relates to number of genes, and the yellow nodes are statistically significant with a p-value $< 0.05$ and false discovery rate $< 0.05$.

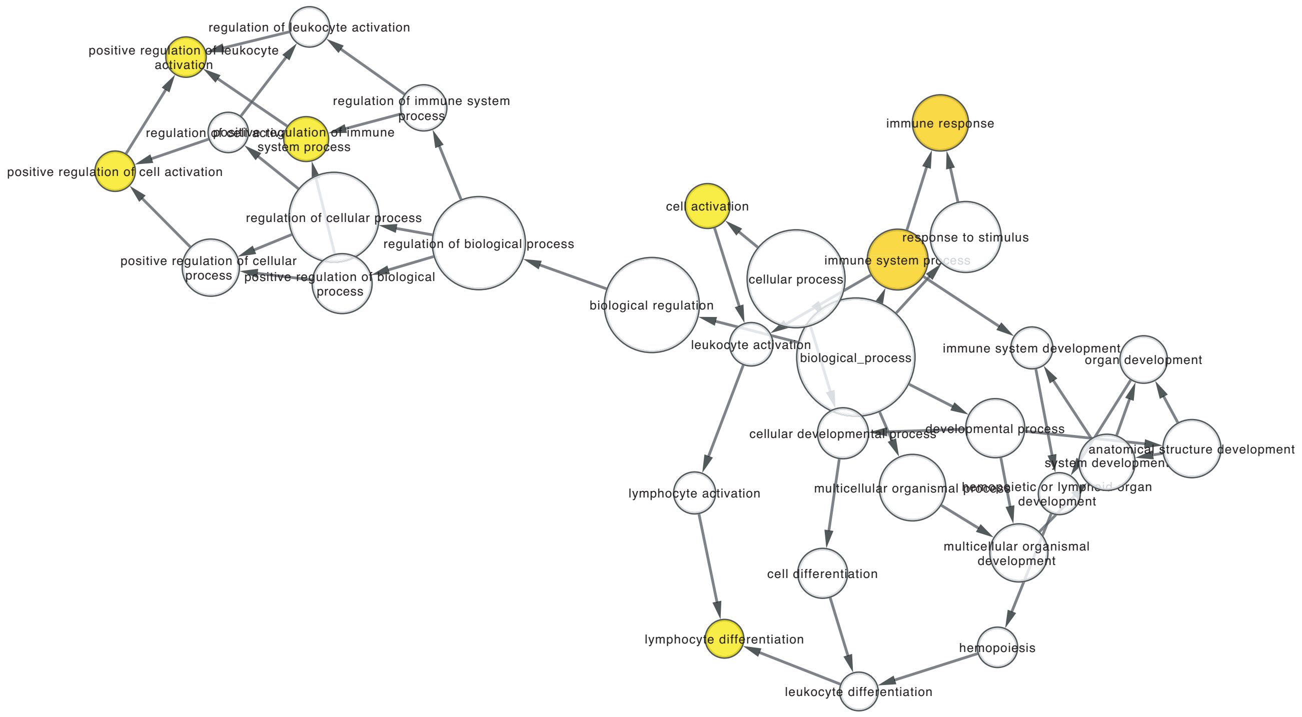


**Figure S5.** Gene Ontology of Biologically Significant Common Genes for the Influenza Infected and Vaccinated Subjects using BiNGO. The node size relates to number of genes, and the yellow nodes are statistically significant with a p-value $< 0.05$ and false discovery rate $< 0.05$.

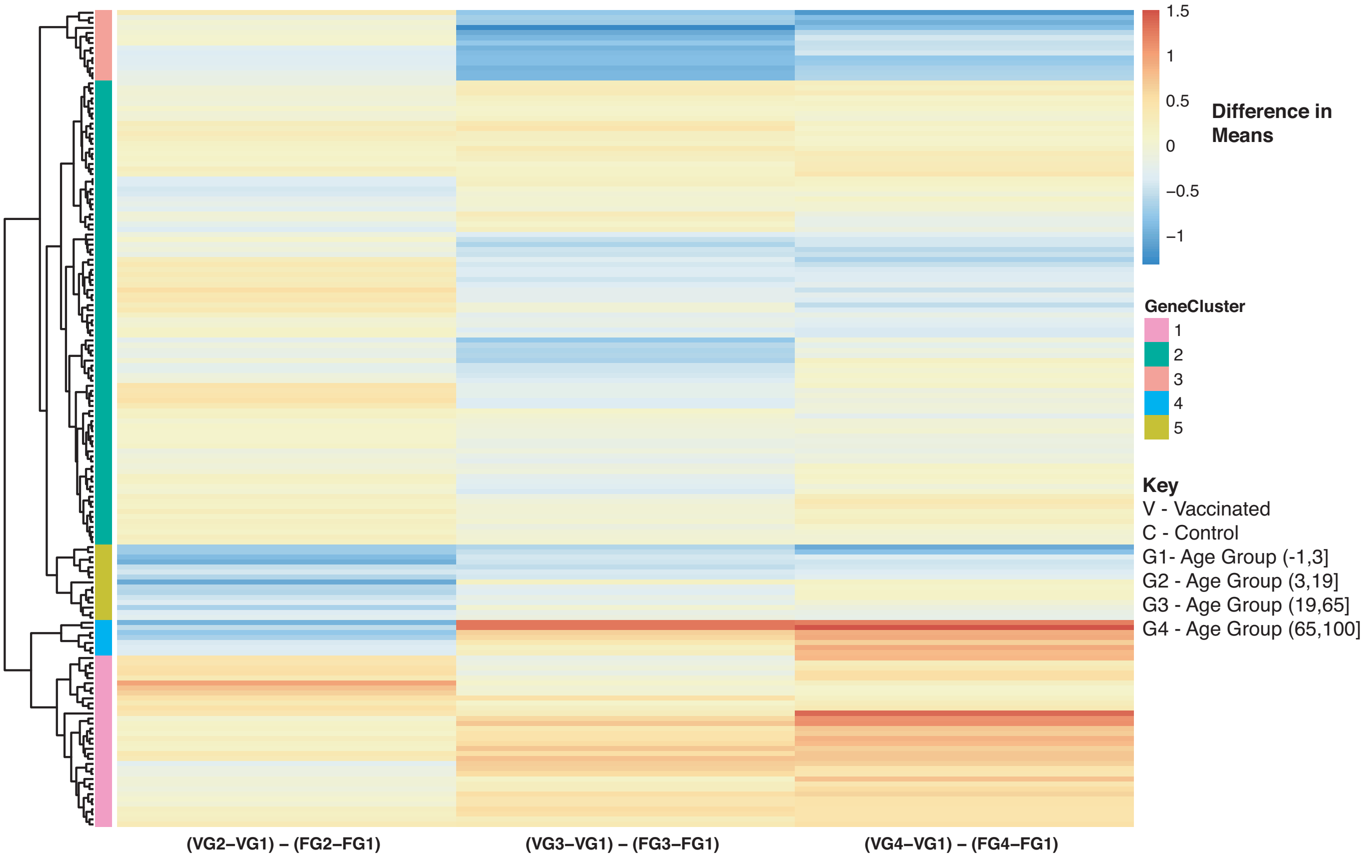


**Figure S6.** Heatmap of Biologically Significant Common Genes for the Influenza Infected and Vaccinated Subjects with an Interaction Between Disease State and Age. Comparison of baseline-adjusted means for influenza vaccinated subjects and influenza infected subjects.

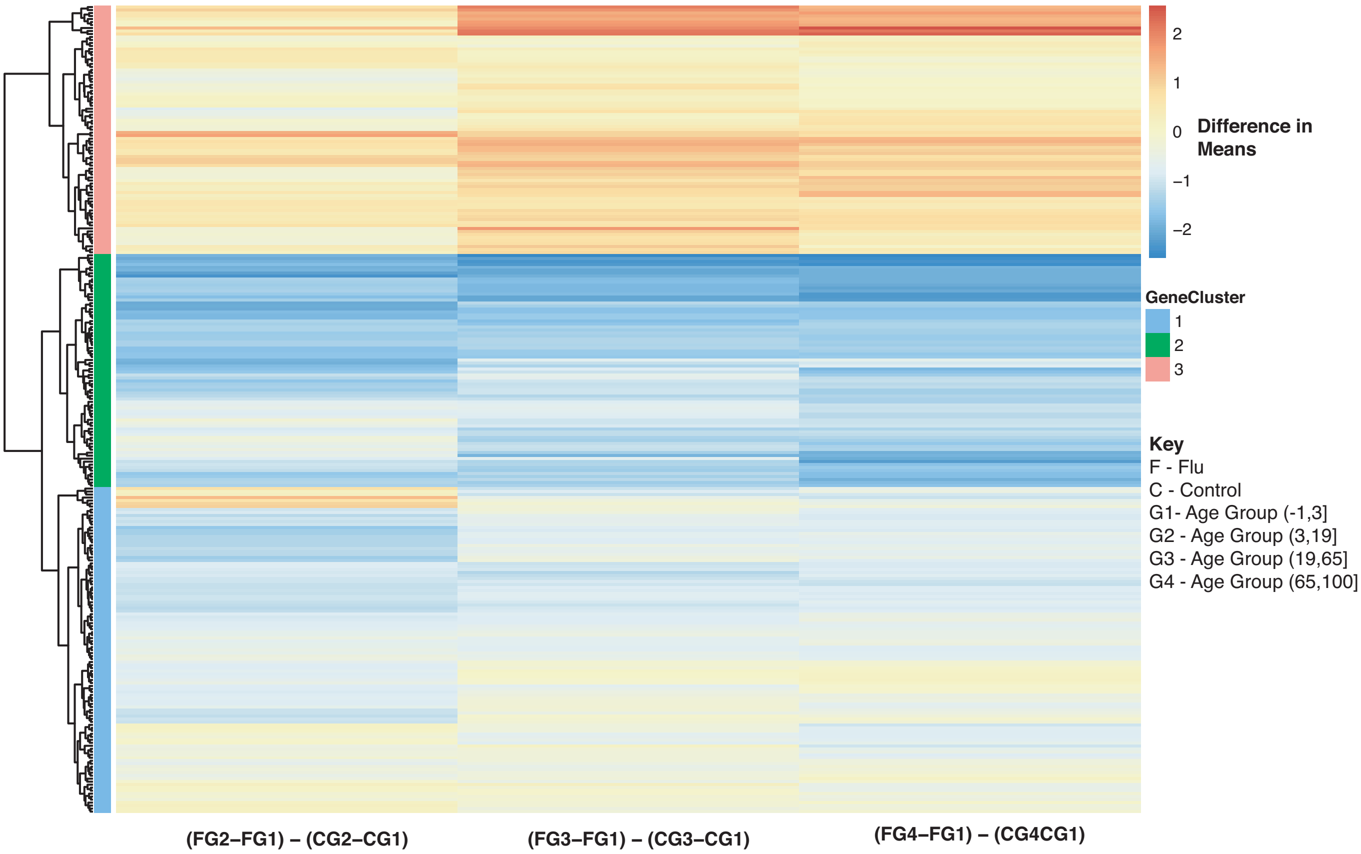


**Figure S7.** Heatmap of Biologically Significant Genes Only in the Influenza Infected Gene List with an Interaction Between Disease State and Age. Comparison of baseline-adjusted means for influenza infected subjects and controls.

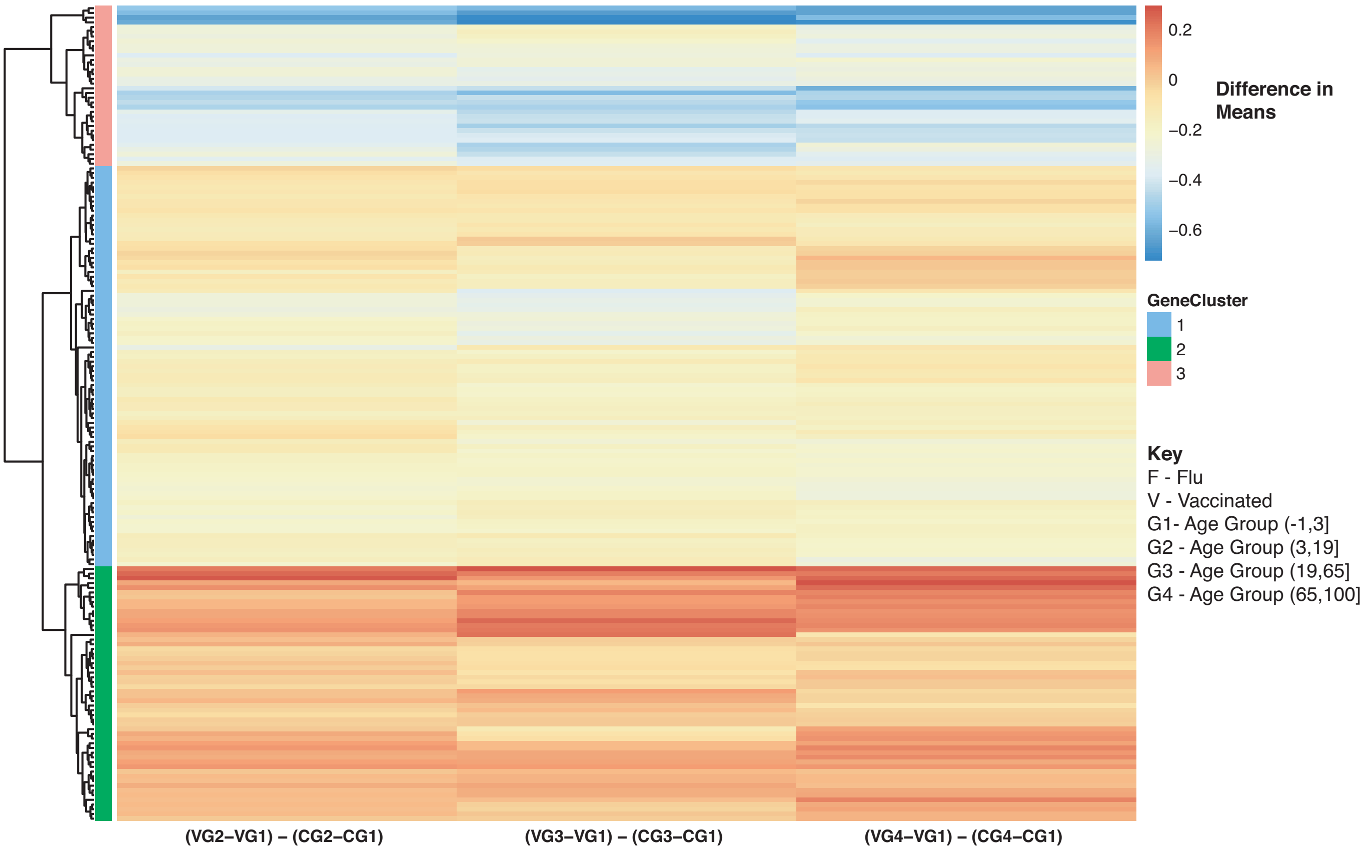


**Figure S8.** Heatmap of Biologically Significant Genes Only in the Influenza Vaccinated Gene List with an Interaction Between Disease State and Age. Comparison of baseline-adjusted means for influenza vaccinated subjects and controls.
